# Reevaluating the *Fusobacterium* Virulence Factor Landscape

**DOI:** 10.1101/534297

**Authors:** Ariana Umana, Blake E. Sanders, Chris C. Yoo, Michael A. Casasanta, Barath Udayasuryan, Scott S. Verbridge, Daniel J. Slade

## Abstract

*Fusobacteríum* are Gram-negative, anaerobic, opportunistic pathogens involved in multiple diseases, including the oral pathogen *Fusobacterium nucleatum* being linked to the progression and severity of colorectal cancer. The global identification of virulence factors in *Fusobacterium* has been greatly hindered by a lack of properly assembled and annotated genomes. Using newly completed genomes from nine strains and seven species of *Fusobacterium*, we report the identification and correction of virulence factors from the Type 5 secreted autotransporter and FadA protein families, with a focus on the genetically tractable strain *F. nucleatum* subsp. *nucleatum* ATCC 23726 and the classic typed strain *F. nucleatum* subsp. *nucleatum* ATCC 25586. Within the autotransporters, we employed protein sequence similarity networks to identify subsets of virulence factors, and show a clear differentiation between the prediction of outer membrane adhesins, serine proteases, and proteins with unknown function. These data have defined protein subsets within the Type 5a effectors that are present in predicted invasive strains but are broadly lacking in passively invading strains; a key phenotype associated with *Fusobacterium* virulence. However, our data shows that prior bioinformatic analysis that predicted species of *Fusobacterium* to be non-¡nvasive can indeed invade human cells, and that pure phylogenetic analysis to determine the virulence within this bacterial genus should be used cautiously and subsequently paired with experiments to validate these hypotheses. In addition, we provide data that show a complex interplay between autotransporters, MORN2 domain containing proteins, and FadA adhesins that we hypothesize synergistically contribute to host cell interactions and invasion. In summary, we report that accurate open reading frame annotations using complete *Fusobacterium* genomes, in combination with experimental validation of invasion, redefines the repertoire of virulence factors that could be contributing to the species specific pathology of multiple *Fusobacterium* induced infections and diseases.

**IMPORTANCE:** *Fusobacterium* are emerging pathogens that contribute to the progression and severity of multiple mammalian and human infectious diseases, including colorectal cancer. Despite a validated connection with disease, a limited number of proteins have been characterized that define a direct molecular mechanism for pathogenesis in a diverse range of host tissue infections. We report a comprehensive examination of virulence associated protein families in multiple *Fusobacterium* species, and show that complete genomes facilitate the correction and identification of multiple, large Type 5a secreted autotransporter genes in previously misannotated or fragmented genomes. In addition, we use protein sequence similarity networks and human cell invasion experiments to show that previously predicted non-invasive strains can indeed enter human cells, and that this is likely due to the expansion of specific virulence proteins that drive *F. nucleatum* infections and disease.

## INTRODUCTION

Bacterial pathogens use a repertoire of diverse virulence proteins to establish infection and confer long-term survival in their respective hosts and environments. A central theme to these virulent phenotypes is the expression of surface exposed and secreted proteins that interact with a variety of macromolecule receptors on host cells (1–4). In addition, these proteins can be deployed by intracellular bacteria within the host cytoplasm or niche-specific vacuole to confer survival, replication, and dissemination. These phenotypes in Gram-negative bacteria are frequently achieved by using large, multi-protein secretion systems, or *nanomachines*, divided into six categories (Type 1-6; T1-6SS)(5). While these are the most common systems to introduce virulence factors into the host or competing bacteria, *Fusobacterium* are unique in that they lack all of the aforementioned multi-protein secretion systems except for the Type 5 secretion system (T5SS)(6). This system is unique in that it is not a large nanomachine, but is divided into five distinct categories (T5a-eSS) that are composed of only one (T5a,c,d,e) or two proteins (T5bSS)(3). These subtypes can be divided into monomeric autotransporters (5a,d)(7, 8), two-partner secretion systems (5b)(9), homo-trimeric autotransporters (5c)(10), and intimins (5e)(11, 12). The majority of characterized autotransporters are large adhesins or proteases of the T5aSS, or homo-trimeric adhesins of the T5cSS that include YadA from *Yersinia pestis (13).* A large scale bioinformatic analysis showed that 100% of Fusobacteria genomes contain T5aSS proteins; the highest percentage in all Gram-negative bacteria tested (14).

*Fusobacterium* are Gram-negative, non-motile, anaerobic bacteria generally isolated from the human oral cavity **(Fig. 1A),** but can also infect other higher mammals including cattle and sheep (15–17). A strong correlation has been established between the presence of *F. nucleatum* in colorectal cancer tumors and a direct induction of increased tumor size, frequency, and stimulation of a pro-inflammatory tumor microenvironment **(Fig. 1B)** (18–20). The interaction of this bacterium with host cells also induces chemoresistance by blocking apoptosis (21), and viable bacteria have been shown to travel within metastatic cells to the liver (22). In addition, increased *F. nucleatum* load within patient sampled tumors correlates with decreased human life expectancy (23).

**FIG 1.**
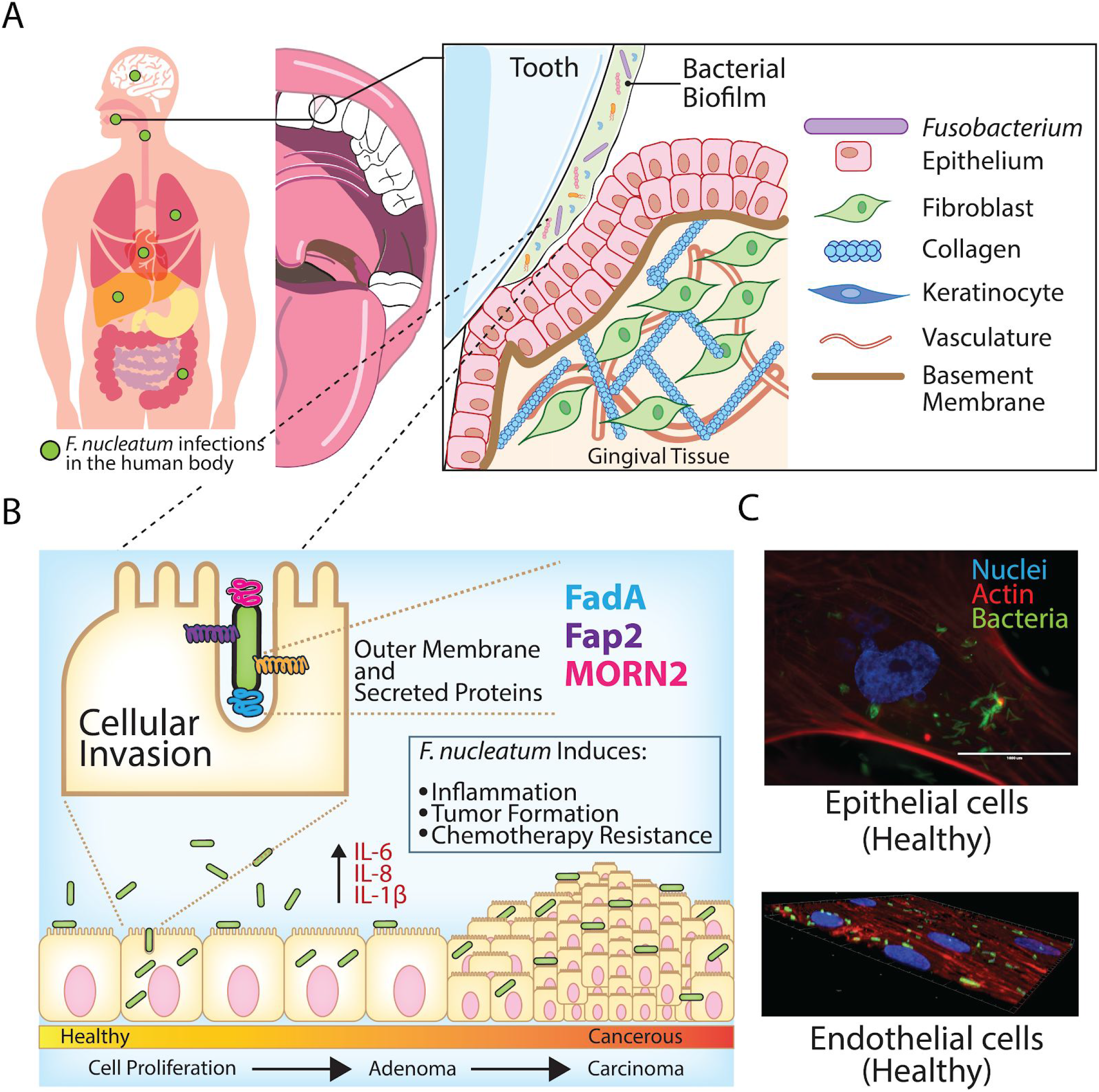
*Fusobacterium* are invasive opportunistic pathogens capable of multi-tissue colonization and infections. (A) Human bodily locations of characterized *Fusobacterium nucleatum* infections, with a focus on the oral environment from which this pathogen disseminates throughout the body. (B) Overview of F. nucleatum cellular invasion, bioinformatically or experimentally characterized proteins that participate in this phenotype, and consequences of infection within localized tissue niches. (C) Fluorescence microscopy showing *F. nucleatum* subsp. *nucleatum* ATCC 23726 is highly invasive in both human epithelial and endothelial cells.

In a recent study, *Fusobacterium* species were bioinformatically divided into actively invading species (*F. nucleatum, F. periodonticum, F. varíum, F. ulcerans*) **(Fig. 1C),** passively invading species that are believed to need a compromised epithelial barrier for cellular entry (*F. necrophorum, F. gonidiaformans*), and those with unknown invasive potential (*F. mortiferum)(24).* Despite the extensive phylogenetic analysis of *Fusobacterium* and our increasing understanding of what is required to be a virulent strain, our knowledge of the specific virulence mechanisms remains limited. For instance, it was previously shown that multiple *F. necrophorum* and *F. mortiferum* strains were significantly more invasive than *F. nucleatum* strains into keratinocytes, which is in direct conflict with bioinformatic reports that place these species in the non-invasive or passively invasive category (24, 25). *F. necrophorum* and *F. mortiferum* are invasive but have not been associated with colorectal cancer, which raises an additional question of how virulence is truly regulated in this disease. Further studies are needed to differentiate the importance of invasion versus only binding to host cell surfaces. For instance, a recent study reported that *F. nucleatum* is able to induce chemoresistance by interacting with surface-exposed toll-like receptors: (21) However, we note that the strain used in this study was highly invasive, yet the amount of intracellular bacteria was not determined. As previously characterized mutants with decreased invasion into human cell lines are also deficient in cellular binding, it could be that there is a complex phenotype that bridges the need for initial host cell docking, and subsequent intracellular modulation of cell signaling.

There is a clear gap in our understanding of invasion and virulence in the genus *Fusobacterium* and the proteins that are involved in driving diverse phenotypes in hosts from humans to cattle and sheep. Most of our knowledge comes from a limited number of *F. nucleatum* strains, and a small sampling of outer membrane adhesins that have been experimentally validated as critical for oral interactions, preterm birth and colorectal cancer. The reason for this lack of molecular studies in *Fusobacterium* is owed to its well known genetic recalcitrance, with only four strains of *F. nucleatum* yielding chromosomal modification (15, 26–28). To aid in our understanding of virulence at the genetic and molecular level, we recently completed nine *Fusobacterium* genomes and created the FusoPortal database that includes detailed genomic and bioinformatic analysis of this emerging pathogen (29, 30). These genomes were used in this study to identify and correct protein families of the autotransporter, FadA, and MORN2 domain containing proteins; all of which are predicted to play key roles in cellular invasion. Among the virulence associated proteins that have been experimentally characterized is Fap2; a large (3786 AA in *F. nucleatum* 23726), dual-function, autotransporter adhesin that binds to the natural killer receptor TIGIT to inhibit tumor cell clearance (30), and initiates host cell docking and altered signaling through the sugar Gal-GalNAc on the surface of colorectal cancer cells (31, 32).

FadA is a small (~125 AA) adhesin that multimerizes on the surface of *F. nucleatum* and has been shown to directly bind to E-Cadherin, where it induces β-catenin signaling in human cancer xenografts in mice (31). For multiple *F. nucleatum* genomes we highlight the identification of two homologues of FadA (FadA, FadA2, FadA3), with multiple identical copies of the FadA3 gene being identified and verified throughout each genome (28). FadA is directly involved in host cell binding and invasion in the strain *F. nucleatum* subsp. *polymorphum* 12230, yet likely due to genetic restraints, has not been characterized in other strains.

In *Fusobacterium necrophorum*, the leukotoxin LktA (*lktBAC* operon) is secreted by the Type 5b Two-Partner secretion system, and has been characterized in cattle and sheep infections causing liver abscesses, spontaneous abortion, and foot rot (*F. necrophorum* subsp. *necrophorum*) (32, 33). In humans, *F. necrophorum* subsp. *Funduliforme is* the predominant subspecies and a native inhabitant of the human oropharynx. This strain is leukotoxin positive and causes infections of the throat and jugular vein in the form of the potentially fatal Lemierre’s Syndrome (26). Despite our knowledge that LktA induces immune cell toxicity, the mechanisms by which these opportunistic subspecies becomes invasive and establish infection in different organisms is poorly understood. To our knowledge, we have sequenced, annotated, and bioinformatically characterized the only two complete *F. necrophorum* genomes in *F. necrophorum* subsp. *necrophorum* ATCC 25286 (34) and *F. necrophorum* subsp. *funduliforme* 1_1_36S (29), which are highlighted through experimental characterization of host cell invasion in this study.

Out of the protein families identified and corrected in this study, the most frequent and extensively enriched are the short, repeated, membrane associated protein domains termed MORN2 (membrane occupation and recognition nexus). This expansion of MORN2 domain proteins is highly specific to *Fusobacterium (24)*, with the exception of multiple genes found in *Helicobacter bilis* which is involved in colitis and hepatitis, as well as infectious abortions in sheep (35, 36). No known function has been assigned to the MORN2 domain containing proteins; however most contain signal sequences allowing for export into the periplasmic space and potential further export to the outer membrane or secretion into the extracellular environment.

With the recent development of a selectable gene knockout system in *F. nucleatum 23726 (27)*, the need to identify accurate gene boundaries and surrounding gene clusters will be critical for studying the virulence proteins used by these bacteria to establish infection. We believe this study will provide a critical tool to drive our improvement of genetic manipulation, therefore allowing previously difficult or impossible cloning and recombinant expression of proteins for the development of antibodies and protein structure-function studies. In addition, these data will ultimately help us answer the question of how a genus of Gram-negative bacteria overcomes a lack of multi-protein secretion machinery which are often necessary for bacterial pathogenesis. In summary, the data in this paper provides a foundation for an increased understanding of how *Fusobacterium* infect a diverse range of host tissues.

## RESULTS

### *Fusobacterium* open reading frame predictions and comparison with previous database annotations

As shown in **Fig. S1,** the previously completed *F. nucleatum* subsp. *nucleatum* ATCC 25586 genome (37) and database depositions of T5aSS proteins were improperly annotated through shortcomings in annotation software, and not errors in the genome. We show that 15 of 15 T5aSS autotransporters, including Fap2 (NP_604343.1: 3165 AA) were previously misannotated (KEGG, Genbank, and UniProt databases) as determined by greatly increased open reading frame length using Prokka (38) and Prodigal (39) and the subsequent identification of the required secretory (SEC) signal sequence for inner membrane translocation. Reannotation of this genome in a previous publication also showed a correction of T5aSS open reading frame annotations (40). To add experimental evidence to our annotations, previous work showed that gene interruption of *radD* in *F. nucleatum* 23726 resulted in a missing protein band as seen by SDS-PAGE at ~370 kDa in *F. nucleatum* 23726, which matches well with our annotated size of this protein at 3461 amino acids. However, the RadD protein was previously annotated as 2143 amino acids in *F. nucleatum* 25586, and our new annotation of 3472 AA matches much better with this experimental data validating a protein of ~370 kDa.

This important initial observation led us to reannotate publicly available *Fusobacterium* genomes to determine if this was a common occurrence. We discovered that multiple genomes suffered from large proteins either being misannotated due to software limitations, or that these large genes spanned the broken boundaries of multi-contig genomes. For example, upon our resequencing and completion of the *F. nucleatum* subsp. *nucleatum* ATCC 23726 genome and comparison with the previous 67 contig draft **(Fig. 2A),** we show that 8 of 15 T5aSS open-reading frames were incorrectly annotated, with four of these being due to contig breaks resulting in fragmented genes. These errors are only partially due to the large gene length (>11kb for fap2), as the other errors were annotation driven **(Fig. 2B).** Many of these proteins are predicted surface adhesins with homology to Fap2; a key protein involved in *F. nucleatum* induced colorectal cancer modulation.

**FIG 2.**
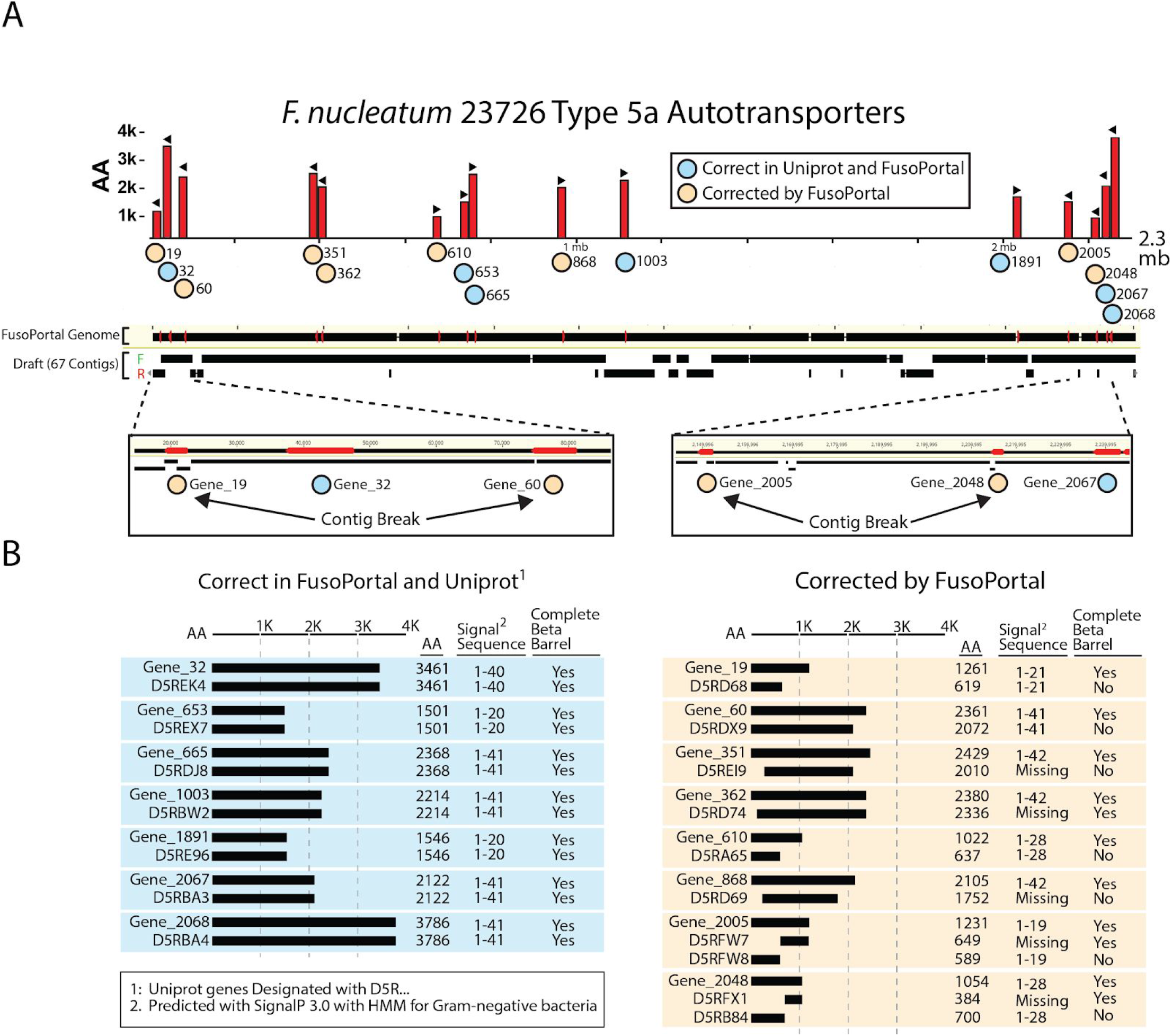
Comparison of T5aSS autotransporter gene annotations from incomplete and complete *F. nucleatum* 23726 genomes. (A) Location (red bar, indicates size in AA) and direction (black arrow) of T5aSS open reading frames in the complete *F. nucleatum* 23726 genome (NCBI: ASM301978v1) from FusoPortal. Contigs from the draft genome (NCBI: ASM17889v1) were aligned on the complete genome and contig breaks that affected gene annotation are highlighted. (B) Autotransporters that were previously correct (light blue) and corrected (tan) are highlighted and show that annotation errors were a combination of N-terminal (signal sequence) and C-terminal (outer membrane embedding β-barrel) truncations.

We subsequently sequenced and completed a total of nine *Fusobacterium* genomes and previously showed that genes larger than 3kb had a high error rate as was validated in this study for the T5aSS proteins (29, 30, 34). A comparison of all annotations of T5a-dSS autotransporters for strains *F. nucleatum* 25586 and *F. nucleatum* 23726 can be found in **Table S1.** In addition, we provide files containing open reading frames for all virulence proteins discussed in this paper in FASTA format (T5SS, FadA, **MORN2)(Text S1),** and **Table S2** contains a list of all open reading frames with InterPro domains and N-terminal signal sequence validation.

### Phylogenetic and protein sequence similarity network analysis of seven *Fusobacterium* species

Full genome phylogenetic analysis of nine *Fusobacterium* genomes spanning seven species **(Fig. 3A)** shows a clear lineage where *F. ulcerans, F. varium*, and *F. mortiferum* are distinct from *F. nucleatum.* Previous analysis had the *F. necrophorum* to *F. nucleatum* phylogenetic relationship more distant than our analysis (23). In the bar graphs in **Fig. 3A,** we highlight groups of virulence proteins as predicted by Hidden Markov Models (HMM), Interpro analysis, and confirmation that there is a Sec dependent signal sequence. For instance, by pure whole genome phylogenetic analysis, previous studies hypothesized that *F. necrophorum* are passive invaders that can not actively induce uptake into host cells. Upon examination of both *F. necrophorum* genomes, we discovered that *F. necrophorum* 25286 has an expansion of *F. nucleatum* like T5aSS autotransporters that share homology with Fap2, while *F. necroporum* 1_1_36S has far fewer T5aSS genes, and none of them cluster with Fap2 and its homologues **(Fig. 3B).** In addition, the non-invasive *F. necrophorum* 1_1_36S nearly lacks MORN2 (Membrane Ontology and Recognition Nexus Type 2) domain containing proteins, which genomically cluster with T5aSS proteins (24), potentially implicating that these proteins may contribute to an invasive phenotype either directly or indirectly. In addition, *F. necrophorum* 1_1_36S does not have any FadA genes, where the invasive strains *F. nucleatum* 23726 and *F. necrophorum* 25286 have five and four, respectively.

**FIG 3.**
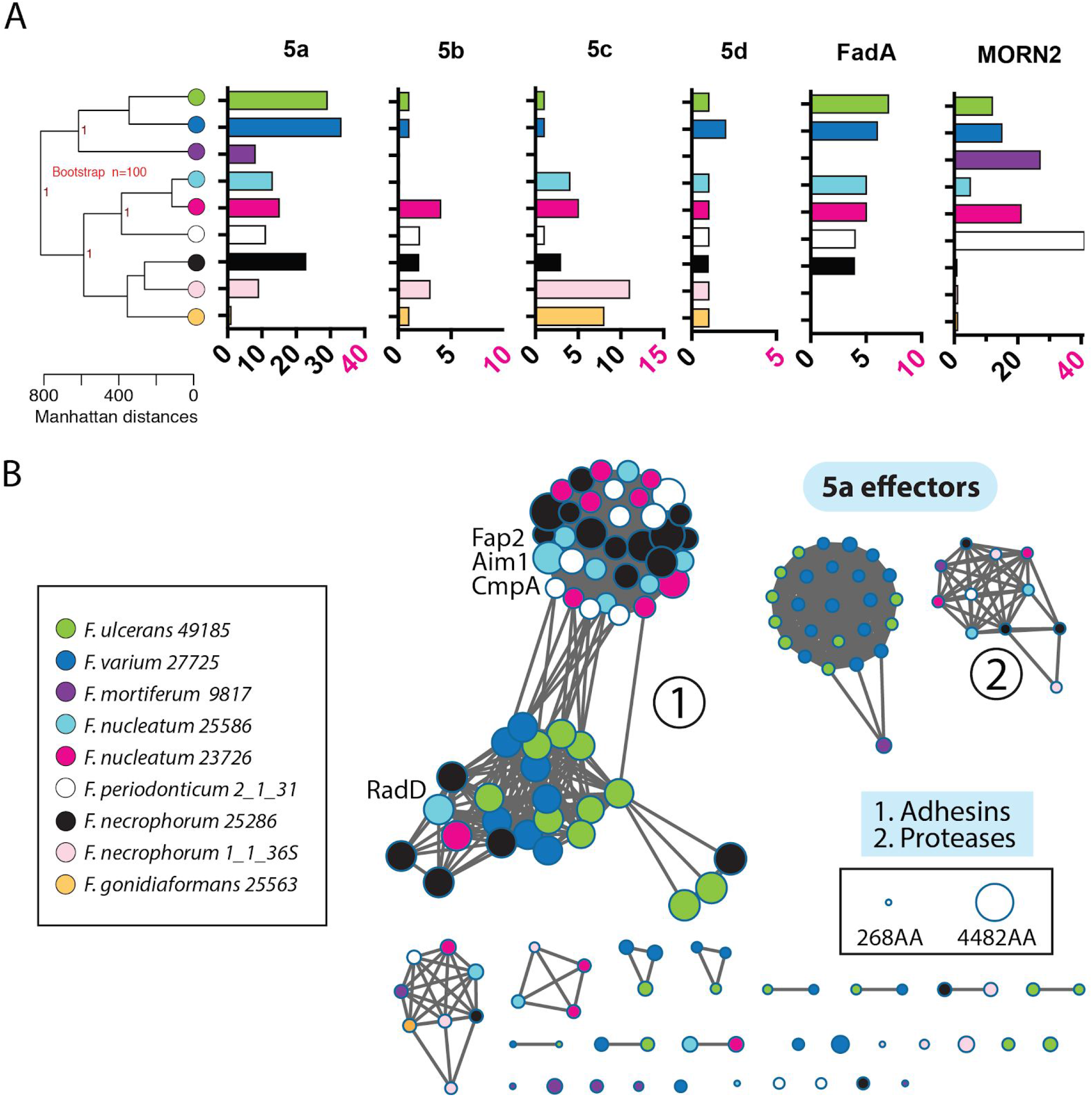
Virulence factor analysis of *Fusobacterium.* (A) Full-genome phylogenetic tree of seven *Fusobacterium* species encompassing nine strains. (B) Sequence similarity networks of all T5aSS autotransporters (e-value 10^−125^) as colored by their designated genomes. Each node represents a single proteins (142 total), and node size represents protein size in amino acids.

To address limitations in predicting the virulence potential of a bacterial strain using whole genomes, we implemented protein sequence similarity networks to determine if any strains and species shared similarity with previously characterized T5aSS virulence proteins from *F. nucleatum* (e.g. Fap2, Aim1)(**Fig. 3B**). To our knowledge, within the *Fusobacterium* autotransporters, no protein sequence similarity networks (SSN) have previously been reported. In addition, we provide a full phylogenetic tree of all 142 T5aSS autotransproters in **Fig S2**. We analyzed all T5aSS autotransporters from nine genomes encompassing seven species of *Fusobacterium*, and show distinct subsets of predicted functions that include surface exposed adhesins, serine proteases, and proteins of unknown function. Wthin the large adhesins, there is a distinct divide between two subsets of T5aSS adhesins at stringent clustering conditions (10^−125^ via EFI-EST)(38). We show that while previous autotransporter analysis broadly clusters RadD with all other Type 5a monomeric autotransporters, this protein associates with a subsection of autotransporters that are more commonly found in multiple copies in *F. varium* and *F. ulcerans.* Worth noting is the absence of Fap2 like adhesins in *F. varium*, yet supernatants from the colonic mucosa of ulcerative colitis patients were able to induce ulcerative colitis in mice (41). When examining the overall virulence landscape between our analyzed stains of *F. nucleatum* and *F. varium*, there doesn’t appear to be much difference when comparing the number of autotransporter, FadA, and MORN2 genes. However, *F. varium* is not associated with colorectal cancer, and we propose that our analysis of differences in outer membrane adhesin subfamilies could be driving the variation in intestinal disorders associated with each species. Since RadD from *F. nucleatum* clusters with the *F. varium* and *F. ulcerans* adhesins, it remains to be determined if RadD plays a critical role in the cellular invasion or the progression of colorectal cancer associated with *F. nucleatum.* More evidence for this comes from the fact that RadD drives interactions with a diverse set of bacteria (39), and is present in a single copy in the two *F. nucleatum* strains analyzed. Oddly, *F. periodonticum* has an expansion of these Fap2 like adhesins, yet has no proteins from the RadD family. To add additional complexity to predicting the function of autotransporters, recent analysis showed that CmpA (Gene_60 in *F. nucleatum* 23726, FN0254), which clusters near the host binding adhesin Fap2, is a crucial adhesin that binds to *Streptococuss gordonii* during oral biofilm formation. These observations may prove crucial in our understanding of protein families that drive interaction networks between the host or competing bacteria in a variety of tissue niches, but provides additional validation that experimental methods are needed to confirm bioinformatic hypotheses.

### Comparison of colorectal cancer cell invasion between

*F. nucleatum* and *F. necrophorum. F. nucleatum* 23726 is a highly invasive strain of *Fusobacterium*, and we show intracellular bacteria inside Ca9-22 cancerous gingival cells in **Fig. 4A.** Because the virulence landscapes between *F. necrophorum* subsp. *necrophorum* 25286, *F. necrophorum* subsp. *funduliforme* 1_1_36S and *F. nucleatum* 23726 were strikingly different (**Fig. 4B),** we tested each strain for the ability to actively invade cancerous colonic epithelial cells (HCT-116). Using imaging flow cytometry, which quantifies the percentage of host cells that are positive for intracellular *Fusobacterium* (fluorescent outer membrane binding lipid)(**Fig. 4C**), we show that *F. nucleatum* 23726 is highly invasive, and *F. necrophorum* 1_1_36S lacks nearly all invasive potential (**Fig. 4D**). This model fits with previous hypotheses that *F. necrophorum* are passive invaders. However, *F. necrophorum* 25286, which has a genetic expansion of *F. nucleatum*-like T5aSS adhesins with homology to Fap2, has significant invasive potential. Our analysis shows that T5aSS effectors are not solely able to predict invasive potential, as *F. necrophorum* 25286 has more genes of this autotransporter protein family than *F. nucleatum* 23726. Upon further evaluation, we hypothesize that autotransporters, FadA, and MORN2 proteins form cooperative networks that individually contribute to host cell docking and invasion. These results show that a more detailed analysis of invasion using additional strains is warranted, and could help in creating predictive models of invasive potential from genome sequences.

**FIG 4.**
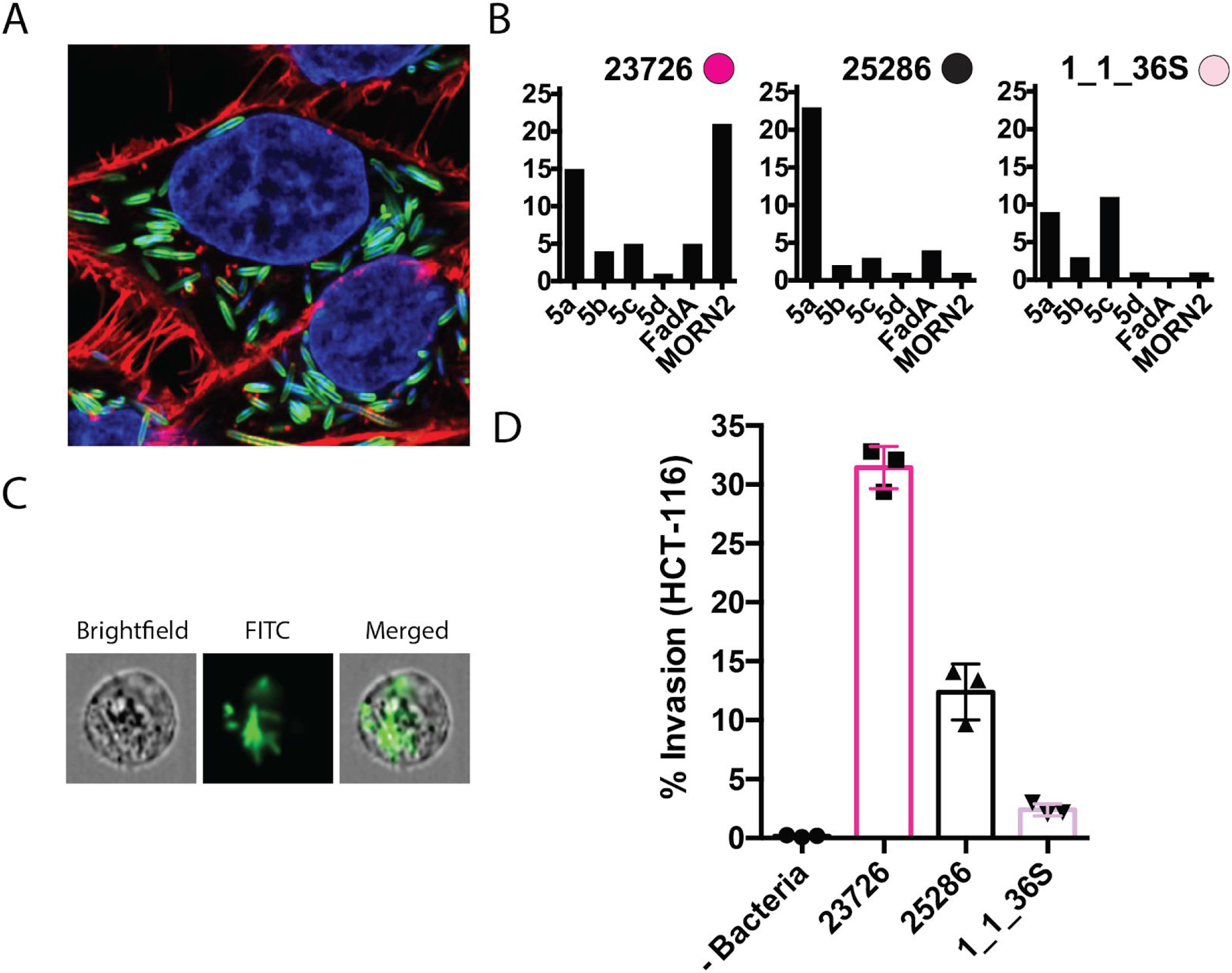
Quantitation of HCT-116 colonocyte invasion by *F. nucleatum* and *F. necrophorum.* (A) Super-resolution imaging of intracellular *F. nucleatum* 23726 in Ca9-22 cancerous gingival epithelial cells. (B) Quantitation of virulence genes in one *F. nucleatum* (23726 and two *F. necrophorum* strains (25286, 1_1_36S). (C) Imaging flow cytometry analysis of Ca9-22 cell invasion using fluorescently labeled *Fusobacterium.* (D) Quantitation of cellular invasion of three *Fusobacterium* strains using imaging flow cytometry.

### A detailed view of T5SS autotransporters in the genetically tractable strain *F. nucleatum* 23726

Autotransporters are Gram-negative specific proteins and constitute the largest family of secreted proteins in bacteria (42). Increasing interest in *F. nucleatum* 23726 has been garnered by several key studies with *fap2* gene interruptions (43, 44), and the recent development of a double-crossover, markerless gene deletion system for this strain (27). In **Fig. 5A,** we highlight the basic structure of the four classes of autotransporters (T5a-dSS) that are found in *Fusobacterium* genomes. *F. nucleatum* 23726 has 25 total autotransporters **(Fig. 5B)** ranging in size up to 3786 AA (Fap2), with the majority falling in the classic monomeric T5aSS proteins. Phylogenetic analysis of the T5aSS autotransporters **(Fig. 5C)** from *F. nucleatum* 23726 fits well with the multi-genome analysis found in **Fig. 3B** and **Fig. S2.** These proteins can bioinformatically be divided into three categories based on predicted function and nearest neighbors: adhesins, serine proteases, and proteins of unknown function. We observed that the previously characterized adhesin RadD from *F. nucleatum* 23726 does not strictly tree near classic adhesins like Fap2 and Aim1, but near the serine proteases that include Fusolisin (FN1426) (39). This could mean that RadD either has an additional uncharacterized function or shares non-enzymatically active motifs with this family. However, a previous study showed that RadD is an arginine-inhibitable adhesin responsible for interspecies bacterial interactions (45). As shown in **Fig. 5D** we have also identified another T5aSS autotransporter adhesin directly upstream in the *F. nucleatum* 23726 genome, of which there has been no characterization. As this gene is not present in the closely related *F. nucleatum* 25586 genome, further research is needed to determine if this protein plays a cooperative role with Fap2 in host cell invasion.

**FIG 5.**
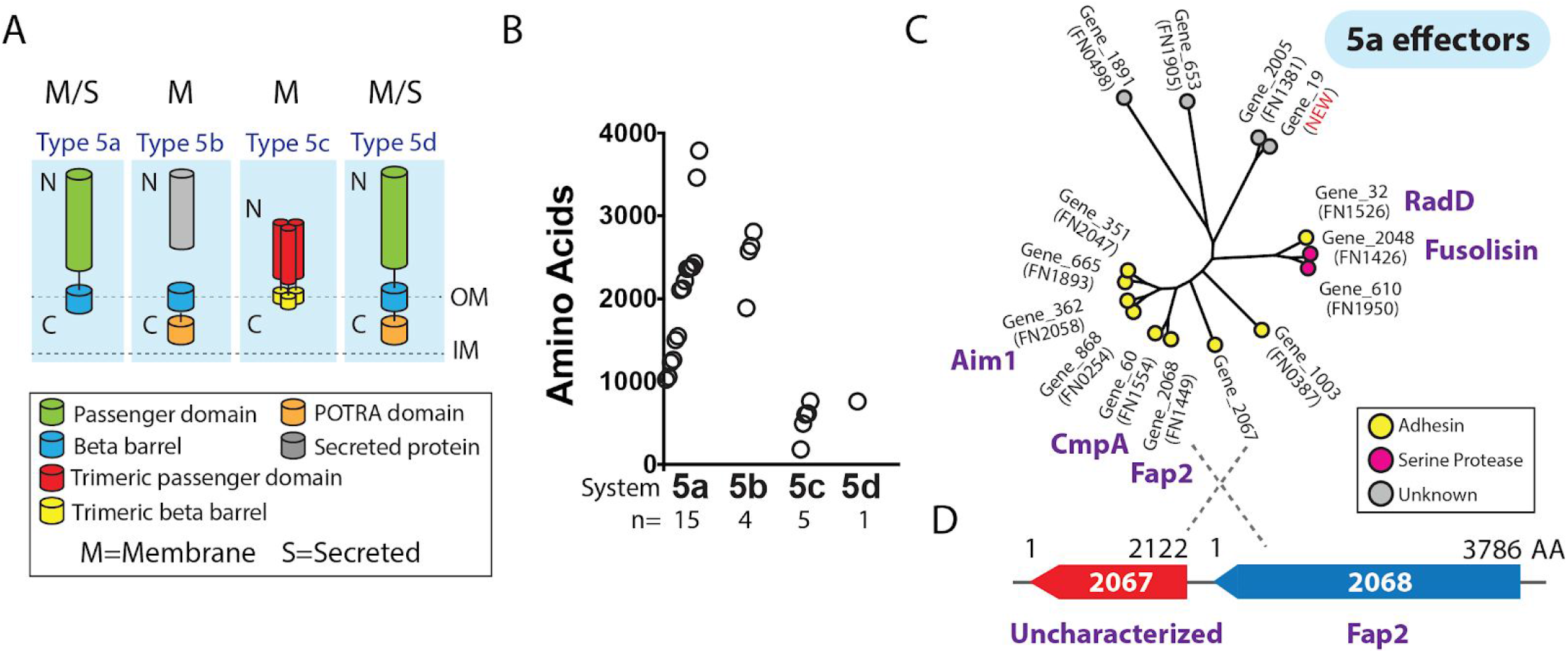
Analysis of T5aSS autotransporters in *F. nucleatum* 23726. (A) Domain structure of the Gram-negative specific, outer membrane bedded or secreted autotransporter protein family. Type 5e (Intimins) is not represented because *Fusobacterium* do not contain these genes. (B) Quantities of each family in *F. nucleatum* 23726, as well as the size in amino acids of each protein. (C) Phylogenetic tree of all 15 T5aSS proteins are divided by node color into predicted protein functions. (D) Highlighting an uncharacterized Fap2 homolog that is just upstream of the well characterized virulence protein Fap2.

Our complete *F. nucleatum* 23726 genome also uncovered a new T5aSS autotransporter (Gene_19) missing in the previous draft assembly. This protein shares strong homology with Gene_2005; previously named FN1381 by the *F. nucleatum* 25586 naming system. Previous bioinformatic analysis of FN1381 in *F. nucleatum* 25586 reported the presence of an ATP- and GTP-binding motif A (P-loop), which currently remains uncharacterized (6). These proteins cluster with multiple autotransporters of unknown function, but could be enzymes based on their phylogenetic locality near the serine proteases. We identified that Gene_19 and Gene_2005 (FN1381) are potential gene duplications that have diverged because the C-terminal ~700 amino acids share 100% sequence identity.

### Discovery and renaming of T5bSS and T5cSS genes in *F. nucleatum* 23726

As shown in **Fig. 6** and **Table S1,** we have identified four full open reading frames for T5bSS effectors (Two-partner secretion – TPS) in *F. nucleatum* 23726, and the full repertoire of these effectors from the nine *Fusobacterium* genomes characterized in this study can be analyzed in a phylogenetic tree in **Fig. S3**. We have renamed the secreted effector genes Type Vb TpsA (*vbaA*, *vbbA*, *vbcA*, and *vbdA*) (**Fig. 6A**). The paired genes encoding for β-barrel translocation proteins were renamed Type Vb TpsB (*vbaB, vbbB, vbcB, vbdB*), and a schematic of TPS architecture is shown in **Fig. 6B.** The previous draft genome of *F. nucleatum* 23726 had the VbaA and VbbA proteins properly annotated, but the *vbcA* gene was fragmented into two open reading frames. *vbdA* is a newly identified gene encoding for a 2634 AA protein. These proteins have not been analyzed in any *Fusobacterium* studies, but other pathogenic bacteria have homologous secreted effectors that contain multiple hemagglutinin domains and function as cytolysins, hemolysins, adhesins, and proteins that initiate contact-dependent growth inhibition (CDI) to fight off neighboring bacteria (46, 47). For examples, a T5bSS filamentous haemagglutinin in *Bordetella pertussis* serves as an adhesin and is essential for colonization of tissues (48). Furthermore, ShlA in *Serratia marcescens* plays a cytotoxic role by contributing to colonization of tissues (49). Thus, while it remains possible that Type Vb autotransporters play a role in tissue colonization, they could alternatively be involved in survival and bacterial competition in oral and colorectal niche environments.

**FIG 6.**
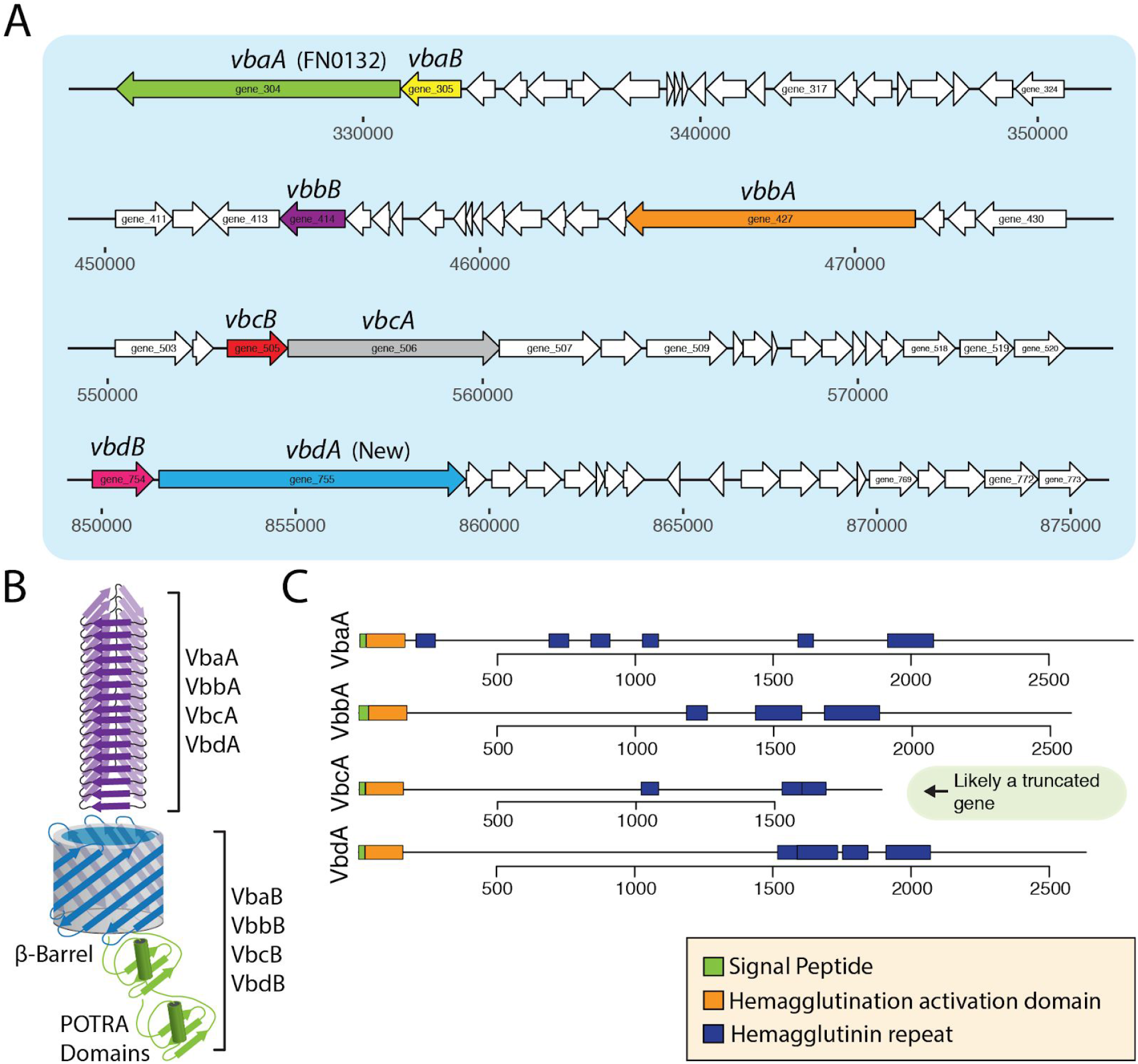
Type 5b secreted proteins (T5bSS) in *F. nucleatum* 23726. (A) Genomic location and open reading frame sizes for four T5bSS effectors with their respective β-barrel translocation proteins. (B) Schematic of the secreted VbaA, VbbA, VbcA, VbdA effector proteins and their respective outer membrane translocation proteins VbaB, VbbB, VbcB, VbdB. (C) Predicted domains of each secreted protein.

In the strain *F. nucleatum* 25586, a previous study revealed that the N-terminal Sec signal sequence was out of frame in T5bSS β-barrel translocation protein, but the presence of a poly-A stretch that bridged this region in the T5bSS β-barrel translocation proteins led the authors to propose that slipped-strand translation could occur during phase variation, therefore translating these signal sequence lacking open reading frames into fully functional proteins capable of translocating a T5bSS effector. Our analysis also identified β-barrel translocation proteins without signal sequences in *F. nucleatum* 25586. However, in the closely related *F. nucleatum* 23726, all open reading frames for the secreted protein and translocation machinery are in frame and are predicted to allow for protein translation. Because of this discrepancy, all *F. nucleatum* 25586 T5bSS systems don’t appear functional, and therefore we haven’t included them in our analysis until biochemical characterization proves the slipped-strand translation and expression of these proteins. We have highlighted all potential *F. nucleatum* 25586 T5bSS genes in **Table S1** and **Table S2,** but note these genes suffer from shifted or in some cases split reading frames.

We report the presence of five T5cSS trimeric autotransporter adhesin (TAA) genes in *F. nucleatum* 23726 as shown in **Fig. 7a,** and we have renamed them *Fusobacterium* Type Vc proteins (*fvcA, fvcB, fvcC, fvcD*, and *fvcE).* Three of the five open reading frames were previously misannotated, with FvcC being the most extreme (new: 615 AA, old: 456 AA). While no TAAs have been studied in *Fusobacterium*, this protein family consists of important virulence factors in other pathogenic Gram-negative bacteria, including YadA from *Yersinia pestis* and *pseudotuberculosis (13, 50)*, Hia from *Haemophilus influenzae (51)*, and SadA of *Salmonella enterica* (52, 53). TAAs form long fibrous proteins that can extend for more than 100 nm from the surface of the bacteria, thereby presenting the adhesive head domains to dock with host cells. YadA binds to the human extracellular matrix proteins fibronectin and collagen, drives invasion into epithelial and phagocytic cells, and inhibits activation of the serum complement (54). In addition to the classic head-stalk-anchor architecture **(Fig. 7B),** several other domains have been characterized that differentiate the role of TAAs (53). As shown in **Fig. 7C,** TAAs can have multiple head domains, and the stalk domains are α-helical coiled-coils. FvcE, differs in that it is quite small at 181 AA, and only contains the required Sec signal sequence and anchor domain. We hypothesized that FvcE could be a non-functioning T5cSS adhesin.

**FIG 7.**
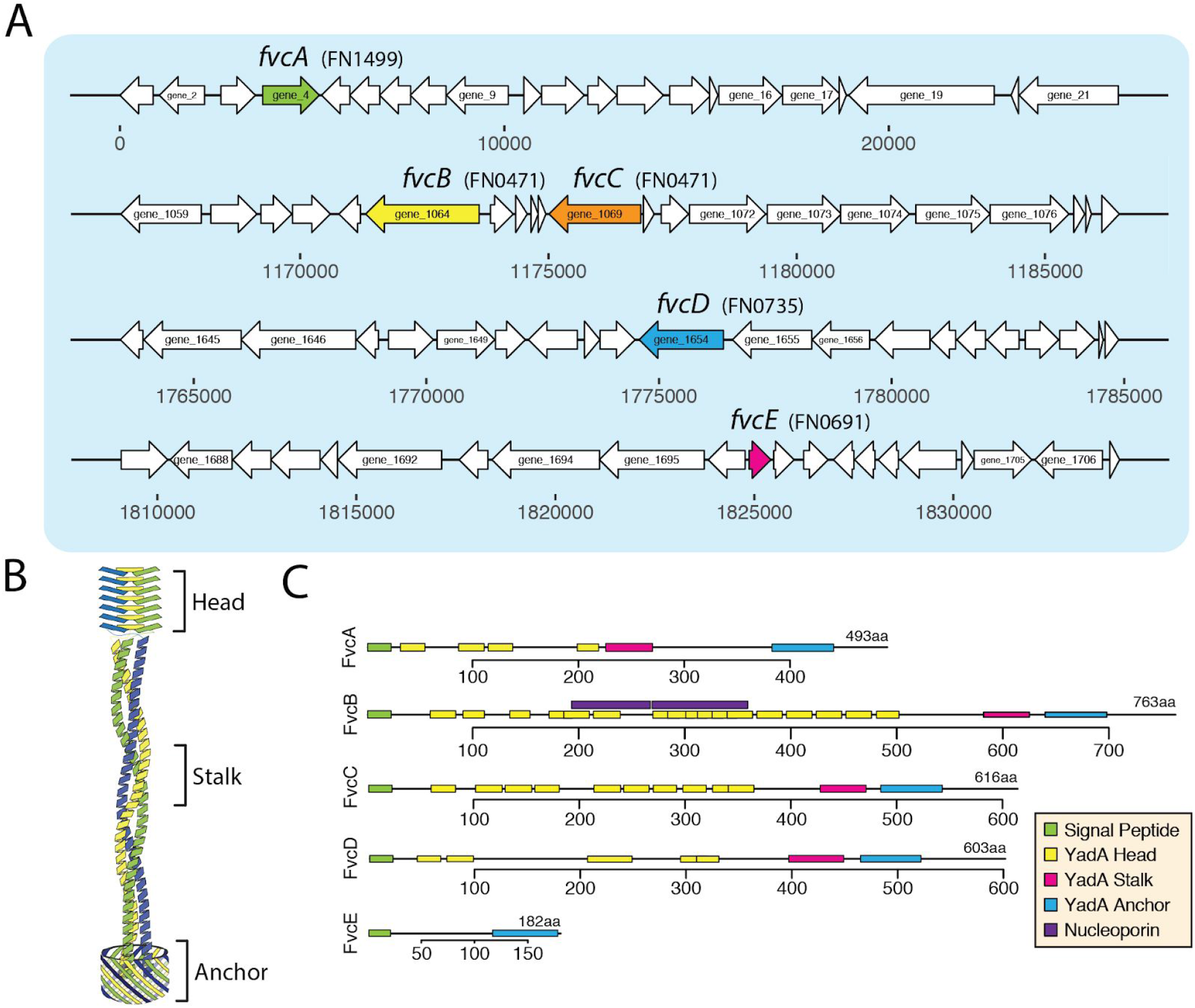
Type 5c secreted trimeric autotransporter proteins (T5cSS) in *F. nucleatum* 23726. (A) Genomic location and open reading frame sizes for five T5cSS trimeric autotransporter proteins. (B) Schematic of the head stalk, and anchor domains. Multiple head domains can be present in one protein. (C) Predicted domains of each each secreted protein.

Previous bioinformatic analysis implicated that a genetic expansion in T5cSS autotransporters in less invasive strains indicates that these proteins are likely not used by *Fusobacterium* for cellular invasion, despite evidence of the TAAs being critical for virulence in multiple human pathogenic Gram-negative bacteria. In **Fig. S4**, phylogenetic analysis of TAAs from nine *Fusobacterium* genomes shows that proteins from highly invasive *F. nucleatum* strains cluster away from those of strains predicted to be less invasive. We propose that TAAs from pathogenic *F. nucleatum* could be playing a role in cellular invasion, and that genetic and biochemical studies need to complement one another before we discount the TAAs as unimportant for *Fusobacterium* virulence.

### Analysis of the FadA protein family

In addition to the Type 5 autotransporters, this current study highlights the unique FadA-like adhesins of *Fusobacterium* **(Fig. 8A-B).** Genetic and biochemical studies of FadA in *F. nucleatum* 12230 showed that this small (~125 AA) adhesin multimerizes on the bacterial surface and subsequently binds to E-cadherin to modulate endothelial barrier permeability, signaling, and inflammatory responses in models of human cancer (55) This implicates FadA in the entry and exit of blood vessels to translocate *F. nucleatum* to the fetal-placental unit where multiple studies support a role for this bacterium in preterm birth (56, 57). Our phylogenetic data **(Fig. S6)** show that multiple ŕađ4-like adhesin genes are present across *Fusobacterium* species, where invasive strains possess a higher number and diversity of these proteins. By contrast, the passive invaders *F. necrophorum* 1_1_36S and *F. gonadioformans* 25563 lack the entire FadA protein family. This newly found FadA profile of *F. nucleatum* warrants further investigation into the invasive potential and function of these paralogs which we have named FadA, FadA2, FadA3a, FadA3b, FadA3c; with the FadA3 proteins being complete gene triplications that are 100% identical at the amino acid level **(Fig. 8C).** The FadA3a-c genes were also found in *F. nucleatum* 25586, as shown by phylogenetic analysis in **Fig. S6,** but were not found in triplicate in the other seven *Fusobacterium* strains analyzed. We validated that these genes were in fact correct annotations and not genome errors using PCR with primers designed for unique upstream regions of each gene in *F. nucleatum* 23726 **(Fig. 8D, Fig. S7)**

**Fig 8.**
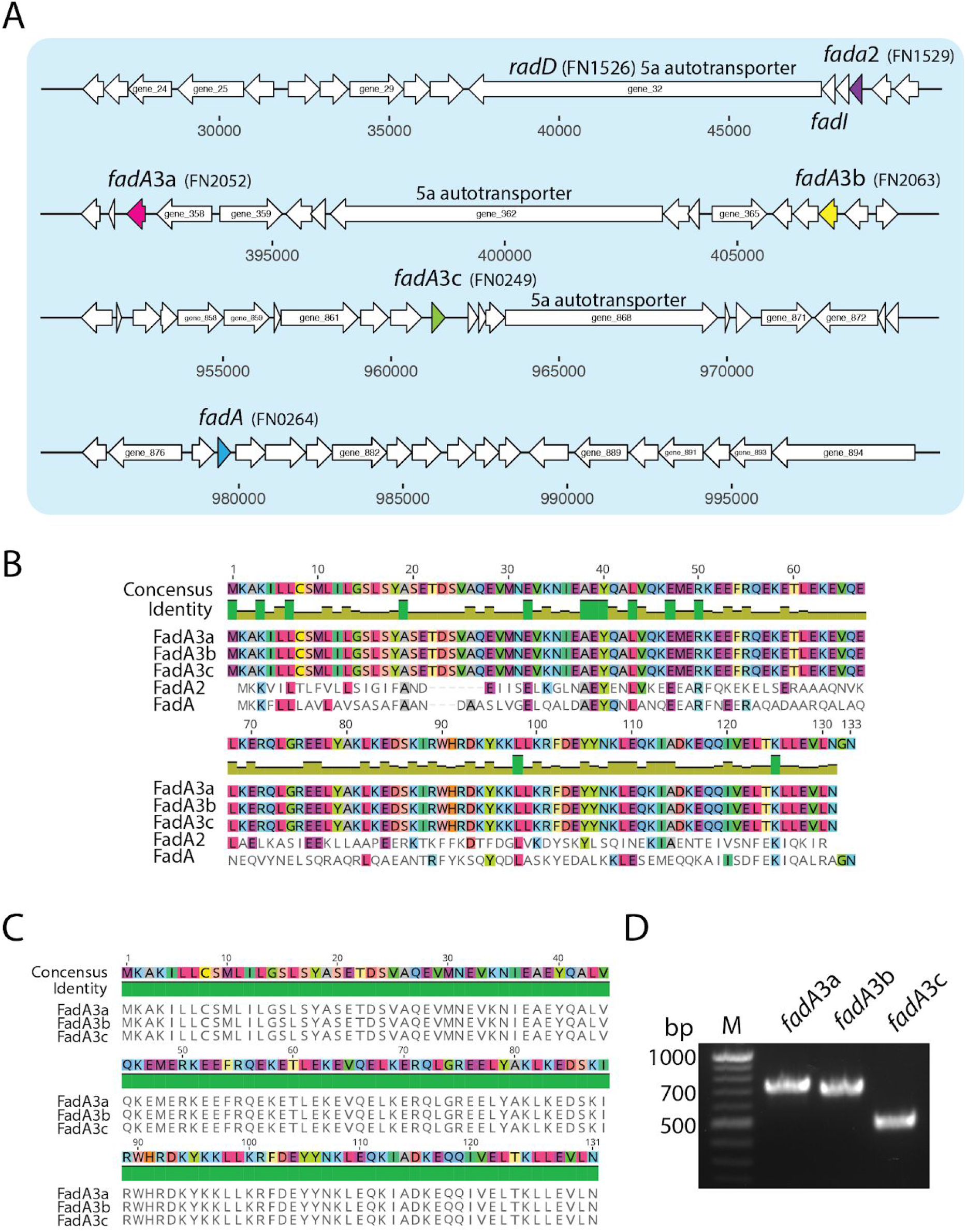
FadA family proteins in *F. nucleatum* 23726. (A) Genomic location and open reading frame sizes for five FadA family proteins. (B) CLUSTAL alignment of FadA proteins from *F. nucleatum* 23726. (C) CLUSTAL alignment of FadA3a, FadA3b, FadA3c shows 100% sequence identity at the amino acid level. (D) PCR to confirm the presence of the *fadA3a, fadA3b, fadA3c* genes at distinct locations in the genome of *F. nucleatum* 23726.

### Towards understanding the role of MORN2 domain proteins in *Fusobacterium* virulence

Genomic studies have greatly detailed the expansion of MORN2 domain-containing proteins in *Fusobacterium*, with the greatest enrichment in the bioinformatically predicted active invaders (24). Despite a great interest in their role in invasion, to our knowledge, no studies have been reported that show a direct role for this hypothesis. In **Fig. S8** we generated a protein tree for 120 MORN2 domain containing proteins from nine *Fusobacterium* genomes. We show in **Fig. 3a** that *F. periodonticum* contains more than 40 MORN2 domain proteins, which is twice as many as the two validated invasive *F. nucleatum* strains.

MORN2 domain-containing proteins could enhance adhesive and active invasive traits as these genes clustered near, FadA, RadD, and additional T5aSS autotransporter adhesins. Since this study, a role for these proteins in interspecies interactions has been suggested (15), as they are predicted surface-associated proteins and some of them are part of the ‘FusoSecretome’ (58). More recently, MORN2 domain containing proteins were found secreted into *F. nucleatum* outer membrane vesicles (59). In addition, the fact that these proteins possess a YwqK domain suggests that they could be acting as toxin-antitoxin systems for interspecies competition or as a bacterial abortive infection systems that limit viral replication and are activated by phage infection (60).

## DISCUSSION

*Fusobacterium* are opportunistic pathogens that cause diverse infections and show strong correlations with multiple diseases in humans and higher mammals (15). *F. nucleatum* has garnered significant attention as an ‘oncobacterium’ that contributes to the progression and severity of colorectal cancer, and has also been implicated in malignant oral leukoplakia (61) and oral squamous cell carcinoma (62, 63). As an oncomicrobe, *F. nucleatum* infection studies have shown that administration of the antibiotic metronidazole reduces tumor burden in human tumor generated xenografts in mice (22). As antibiotic therapy to treat disease could result in altered gut flora that changes the efficacy of chemotherapy drugs, a new paradigm would be to control infections at the disease site without antibiotics, or the opportunity to block this pathogen from leaving its native human oral cavity. For *F. necrophorum*, mostly of the subspecies *funduliforme*, infections of the jugular vein and subsequent progression to Lemierre’s Syndrome are often fatal in humans. *F. necrophorum* subsp. *necrophorum* is the primary causative agent of bovine hoof rot and liver abscess, causing severe monetary loss in the livestock industry. The differences in *F. nucleatum* and *F. necrophorum* virulence and disease association is not well understood. We highlight that both *F. necrophorum* 25286 and *F. necrophorum* 1_1_36S both encode for the LktA leukotoxin that induces the activation and apoptosis of leukocytes (64, 65). The *lktBAC* operon responsible for the production of this T5bSS secreted toxin is absent in all other strains of *Fusobacterium* analyzed, and could explain the severe abscess phenotype induced by this species.

There have been several bioinformatic studies of *Fusobacterium*, but none of these used databases that were entirely populated with complete genomes. This lack of complete genomes led us to sequence, assembly, annotate, and create the FusoPortal database (29) of *Fusobacterium* genomes to aid in the bioinformatic and molecular experiments found in this study. These data provide highly accurate gene boundaries, and therefore will greatly facilitate the design and production of recombinant proteins for structural and functional studies; an area that is severely lack in the *Fusobacterium* field. Our data confirmed that *Fusobacterium* are unique in that they lack all large protein secretion machinery of the Type 1,2,3,4, and 6 varieties, yet they are still opportunistic pathogens. However, a recent proteomic analysis of *Fusobacterium nucleatum* revealed that secreted outer membrane vesicles consist of proteins from of each of the virulence factor families we analyzed in this study (57).

We show the importance of having accurate and complete genomes in identifying large proteins; the majority being Type 5 secreted proteins that have been characterized as virulence factors in a wide range of Gram-negative bacteria. As we have seen the importance of a single protein in *F. nucleatum* virulence in the outer membrane adhesin Fap2, it will be important to not overlook proteins of this size merely because incomplete genome assembly results in open reading frame fragmentation. A key contribution of this work is the identification of Fap2 homologues in multiple *Fusobacterium* genomes, with a focus that *F. necrophorum* subsp. *necrophorum* 25286 has two close homologues of this adhesin. As this subspecies of *F. necrophorum* is associated with serious livestock infections, this data, combined with the potential to perform genetic manipulation in these strains, could lead to a more effective live, attenuated vaccine for bovine hoof rot and liver abscesses. In addition, now that all large outer membrane autotransporters have been identified in multiple *F. necrophorum* strains, there is an opportunity to develop new protein-based vaccines from this proven immunogenic protein family (pertactin, *Bordetella pertussis*, whooping cough vaccine)(66). In *Orientia tsutsugamushi*, the autotransporter ScaA acts as a critical bacterial adhesion factor. Animal models have shown that anti-ScaA antibodies have proven to be the most promising trial of scrub typhus vaccination (67). Finally, The T5aSS autotransporter Hap from *Haemophilus influenzae* has been studied as a potential vaccine target for the prevention of nontypeable *H. influenzae* disease (68, 69).

While we know that *Fusobacterium* contribute to several diseases, our understanding of the overarching molecular mechanisms driving the virulence of these bacterial species remains limited. We focused our virulence factor analyses on the Type 5 autotransporters, FadA adhesins, and MORN2 domains protein families. We briefly state that to our knowledge, the role of T5bSS (Two-partner secretion), T5cSS (trimeric autotransporters), and MORN2 domain proteins in Fusobacterium have no published functional studies, and therefore represent an exciting future field to pursue. Not analyzed in this study, Fad-I is an outer membrane protein that was shown to induce human beta defensin 2 (hBD-2) through a toll-like receptor mediated host response. The immune modulation induced by Fad-1 in *F. nucleatum* strains 23726 and 25586 was far more potent than that seen in *F. nucleatum* 10953 (70). As with all protein families, this is a great example of how small sequence variations in key proteins could account for altered virulence between phylogenetically similar strains of *Fusobacterium.*

Type 5a secreted autotransporters are virulence factors in a broad range of Gram-negative pathogens but appear to rely heavily on an expansion of T5SS effectors for host colonization and infection. *F. nucleatum* in the oral cavity serves as a bridge for bacteria-bacteria aggregation, and interactions with mammalian cells and inert tooth surfaces within the gingival pocket (71). The T5aSS proteins RadD and Fap2 are critical for inter-species adherence and the overall architecture of multispecies biofilms (43, 45). Therefore, Type 5a autotransporters play key roles in co-aggregation, cell-cell interaction, and biofilm formation during healthy and pathogenic states. Our data suggest that newly identified subsets of T5aSS auotransporters are present in a variety of *Fusobacterium* species, and therefore new genetic and biochemical studies should be designed to characterize these proteins in virulence.

While *Fusobacterium* have been reported as non-motile, we seeked to determine if *F. nucleatum* T5aSS autotranspoters share homology with IcsA from from *Shigella flexneri*, or T5cSS proteins with BimA from *Burkholderia* species, as these proteins localize to a single bacterial pole (old pole) and coordinate intracellular actin based motility (72, 73). Using our custom BLAST server that is built into the FusoPortal database, we show no homology to IcsA and low homology to the membrane associated β-barrel of BimA. These data agree well with a lack of actin based motility reported by all previous studies and our own observation of intracellular *F. nucleatum.* Despite not using actin-based motility for intracellular movement, another intriguing observation was that actin localizes to intracellular *F. nucleatum* in human keratinocytes (74), and that inhibition of new actin synthesis blocked intracellular entry (74). The protein or proteins involved in this direct or indirect actin recruitment have yet to be identified. Determining if intracellular *F. nucleatum* cloak themselves or a host vacuole in actin to evade host clearance through ubiquitination and xenophagy, as seen for *Listeria monocytogenes* and *Chlamydia trachomatis (75, 76)*, could be a key piece in understanding the intracellular persistence and dissemination of this pathogen.

The identification of up to six FadA family proteins in a genome leads us to hypothesize that there is cooperativity among this protein family, and that the FadA homologues, FadA2 and FadA3, could have similar functions as adhesins, but have unidentified host receptor molecules. In *F. nucleatum* 12230, a *ΔfadA* mutant shows greatly reduced proliferation of human cancer cells (HCT-116, HT29), but this pro-carcinogenic phenotype was recovered by *fadA* complementation and the addition of purified active FadA to cell cultures. In addtion, FadA is also important for the colonization of *F. nucleatum* 12230 in mouse placenta (77). With five FadA family proteins in the genetically tractable F. nucleatum 27326, it will be key to delete multiple copies of these proteins to determine if they act synergistically during infection, or if FadA, FadA2, and FadA3 play distinct roles in virulence and colonization in diverse tissue niches including the subgingival microbial community, human placenta, or the colon.

In conclusion, we provide bioinformatic identification and analysis of virulence factors, as well as host-pathogen infection studies to dissect the role of this landscape in cellular invasion. We hypothesize that autotransporters, FadA, and MORN2 proteins synergistically form a host cell docking and invasion network that confer the host, tissue, and disease mechanism of the diverse range of *Fusobacterium* species. These results show that a more detailed analysis of invasion using additional strains is warranted, and could help in creating more accurate predictive models of invasive potential from genome sequences. This study will benefit from future work that expands upon our nine genome analysis to include additional *Fusobacterium* species and clinical strains, which will ultimately lead to a deeper understanding of individual proteins in disease. It is our hope that these genomes, bioinformatic analyses, and *Fusobacterium* invasion studies spark the discovery of new virulence factors and drive studies that further our understanding of virulence mechanisms at the molecular level in the diverse bacterial genus of *Fusobacterium.*

## MATERIALS AND METHODS

### Use of genomic information for bioinformatic analysis of nine *Fusobacterium* strains

*Fusobacterium* genomes and all associated data used for this study can be accessed under the NCBI BioProjects PRJNA433545 and PRJNA513186. Annotations for all genes were performed with Prodigal (39) or Prokka (38), and can be found on the FusoPortal website (http://fusoportal.org) or our Open Science Framework data repository (http://osf.io/2c8pv). In addition, NCBI annotated each genome using their Prokaryotic Genome Annotation Pipeline. Signal peptide prediction was done with SignalP 3.0 and the HMM option (78). Protein functions were predicted using Interpro analysis (79) within Blast2GO 5.2.5 software (80), and predictions for all virulence proteins used in this study can be found in **Table S2.**

### Protein sequence similarity networks (SSN)

Networks were created by identifying all protein families of interest using the Interpro server within Blast2Go software. Protein hits were manually checked for accuracy and the presence of a Sec signal sequence. Qualifying proteins were then analyzed using the all-vs-all BLAST feature within the EFI-EST server (81), followed by cluster mapping in Cytoscape 3.0 (82). All raw files for proteins in fasta format, as well as Cytoscape networks, are posted in our Open Science Framework repository.

### Phylogenetic analysis of *Fusobacterium* genomes

Full genome phylogenetic analysis of nine *Fusobacterium* genomes **(Fig. 3A)** was performed using the MicroPan (83) R-package of pan-genome analysis. All protein open reading frames for each genome were included in the analysis. Nodes for each protein were manually colored in Adobe Illustrator to match the genome key depicted in **Fig. 3B.**

### Phylogenetic analysis of virulence factors

Protein trees **(Fig. S2–6, S8)** were built within Geneious 9.02 using the Geneious Tree Builder. Pairwise alignment was run using a Blosum62 cost matrix and neighbor-joining trees were built using a Jukes-Cantor genetic distance model. Nodes for each protein were manually colored in Adobe Illustrator to match the genome key depicted in **Fig. 3B.**

### Cellular infections of *F. nucleatum* in epithelial and endothelial cells

A single colony was used to start overnight cultures of *F. nucleatum* 23726, *F. necrophorum* 25286, or *F. necrophorum* 1_1_36S. Stationary phase cultures were back-diluted to OD_600_ = 0.1 in 2 mLs of Columbia Broth supplemented with hemin and vitamin K (CBHK) and grown to exponential phase (OD_600_ = 0.4). *Fusobacterium* cultures were then transferred to a 1.5 mL conical tube and centrifuged at 5,000 rcf for 3 minutes to pellet bacterial cells. CBHK was removed and the bacterial cell pellet was resuspended in 100 uL of PBS, pH 7.4, and incubated in 500 ng FM 1-43FX membrane probe for 5 minutes. Bacteria were centrifuged at 5,000 rcf for 3 minutes, and the cell pellet was washed with PBS, centrifuged at 5,000 rcf for 3 minutes, and the cell pellet was resuspended in 1 mL of PBS pH 7.4. Next, *Fusobacterium* were added (MOI 10:1) to the media of confluent human cell monolayer cultures in 6 well plates (for imaging flow cytometry) or #1.5 coverslip bottom plates (fluorescence microscopy), and incubated for 1 hour at 37 °C in 5% CO_2_. Following infection, epithelial cells were washed two times with cell culture media and the extracellular bacteria were killed by incubating epithelial cells with cell culture media supplemented with penicillin/streptomycin for 30 minutes. Post infection, cells were either analyzed by fluorescence microscopy or analyzed by imaging flow cytometry as described in the following sections.

### Fluorescence microscopy of intracellular *F. nucleatum* in FHC colonocytes and Ca9-22 gingival cells

In **Fig. 1C** (top panel), a confluent monolayer of FHC healthy colonocytes (ATCC CRL-1381) were grown in DMEM:F12 medium supplemented with 10% FBS (Atlanta Biologicals) in #1.5 coverslip glass bottom plates. Cells were infected in antibiotic free DMEM:F12 media with a 10:1 MOI of *F. nucleatum* 23726 labeled with the fluorescent lipid FM 1-43FX were for one hour at 37 °C in 5% CO_2_. Extracellular bacteria were then killed with DMEM:F12 supplemented with 100 I.U./mL penicillin and 100 (μg/ml_) streptomycin. After washing with media, cells were fixed in well with PBS 3.7% paraformaldehyde, and permeabilized with 1% Triton X-100. Cells infected with *F. nucleatum* 23726 were then stained with 1.0 μg/ml DAPI (Invitrogen) to visualize DNA and Texas Red-X phalloidin for F-actin filaments (Thermo Fisher). Images were acquired using a63Xoil objective on an EVOS FL fluorescent microscope.

In **Fig. 4A,** Ca9-22 cancerous gingival cells were used for invasion with *F. nucleatum* 23726. Growth and infection conditions were the same as for the HCT-116 cells above with the exception that the culture media was MEM/EBSS with 10% FBS. Images were acquired using a 60X objective on a Zeiss 880 Airyscan equipped fluorescent microscope.

### Fluorescence microscopy of intracellular *F. nucleatum* HVMEC endothelial cells

In **Fig. 1C** (bottom panel), HVMEC Adult Dermal Cells (Lonza) were cultured in a custom-built 3D printed chamber where a #1.5 coverslip was adhered to the bottom of the printed well using biocompatible polydimethylsiloxane ((PDMS) SYLGARD 184 Silicone Elastomer Kit). The CAD model of the micro-chamber was designed using Autodesk Inventor Professional 2018. The chamber was printed with PLA using MakerBot Ultimaker 2.. Additionally, cell attachment factor (Cell Systems) and a thin lining of collagen (5mg/ml_) were coated on the coverslip before seeding the host cells within the wells. HVMEC cells were grown in EGM MV-2 (Lonza) media at 37 °C in 5% CO_2_. Cells were infected in antibiotic free DMEM:F12 media with a 10:1 MOI of F. nucleatum 23726 labeled with the fluorescent lipid FM 1-43FX were for one hour at 37 °C in 5% CO_2_. Extracellular bacteria were then killed with DMEM:F12 supplemented with 100 I.U./mL penicillin and 100 (μg/mL) streptomycin. After washing with media, cells were fixed in well with a 10% formalin solution. The fixed cells were washed with phosphate buffer solution (PBS) and blocked with a solution of PBS-X containing 10 mL PBS, 5μL Triton-X, and 200 mg Bovine Serum Albumin (Fisher BioReagents). After rinsing with PBS, the cells were stained with Alexa Fluor 568 phalloidin (Life Technologies) for F-actin filaments (1:100 Dilution) and 1.0 μg/ml DAPI (Invitrogen) for the nucleus. Imaging was performed using a Zeiss LSM 800 Confocal Microscope. Representative images and z-stacks were obtained using a Zeiss 63x 1.8 NA oil immersion lens with 4x averaging.

### Imaging Flow Cytometry and quantitation of *Fusobacterium* invasion into HCT-116 cells

At the end of two hour *Fusobacterium* infections, adherent HCT-116 cancerous human colonocytes were gently removed from tissue culture treated plates by incubating with 0.05% trypsin for 5 minutes and subsequently transferred to a 1.5 mL conical tube. The trypsin was then quenched with 1 mL of cell culture media supplemented with penicillin/streptomycin and 10% FBS Cells were centrifuged at 1,000 g for 3 minutes. Pelleted epithelial cells were then fixed by incubating in 3.2% paraformaldehyde for 15 minutes. Following incubation, the cells were centrifuged at 1,000 g for 3 minutes, and pelleted cells were washed one time with 1 mL of PBS pH 7.4. Cells were then centrifuged at 1,000 g for 3 minutes and resuspended in 30 μL of PBS pH 7.4 and visualized using AMNIS ImageStream Mark II instrument. For each data set in triplicate, > 1000 individual cells were selected by selective gating applied to all data sets, and % invasion was determined by dividing the number of FITC channel positive cells by the total number of cells.

### PCR to validate the presence of three separate genes for *fadA3a-c* in *F. nucleatum* 23726

Genomic DNA was extracted from exponential phase *F. nucleatum* 23726. Primers as shown in **Fig. S7** are prDJSVT1006-1009. The following 5’-3’ correspond to each: prDJSVT1006; 5’ CT AAACT CTTCCTTT CTTTCCATTTCC 3’, prDJSVT1007; 5’ CCT AT CAGGAAGT AT GAT AACTTT AACAAG 3’, prDJSVT1008; 5’ CCTATACCCTAGCAAACAATAACAAAGTC 3’, prDJSVT1009; 5’ GAACATGAAATAAGACAAATATTTTAAGGATG 3’. PCR was run using Q5 polymerase (NEB) and a 50-62 °C annealing temperature gradient. Products were visualized using 1% agarose gels stained with ethidium bromide.

### Data availability

All genomic data used for protein annotation can be found on NCBI under BioProjects PRJNA433545 and PRJNA513186. All associated raw data outside of supplemental materials, which includes the T5aSS autotransporer protein similarity network can be found on our our Open Science Framework data repository (https://osf.io/vs3fd/?viewOnlv=1752886f47234a6e900f75b73b72fe56)

## ACKNOWLEDGMENTS

This article has been supported by the National Science Foundation Career Award CBET-1652112 (Verbridge), a Commonwealth Health Research Board Award 208-10-18 (Slade), and the USDA National Institute of Food and Agriculture (Slade). We would like to thank the following for technical assistance with experiments: Kristi Decourcy from the Fralin Life Sciences for assistance with microscopy; Melissa Makris from the Virginia-Maryland School of Veterinary Medicine for imaging flow cytometry. We thank Virginia Tech’s Open Access Subvention Fund for publication funding.

## SUPPLEMENTAL MATERIAL

### SUPPLEMENTAL FIGURES. Umana et al. 2019

**FIG S1.**
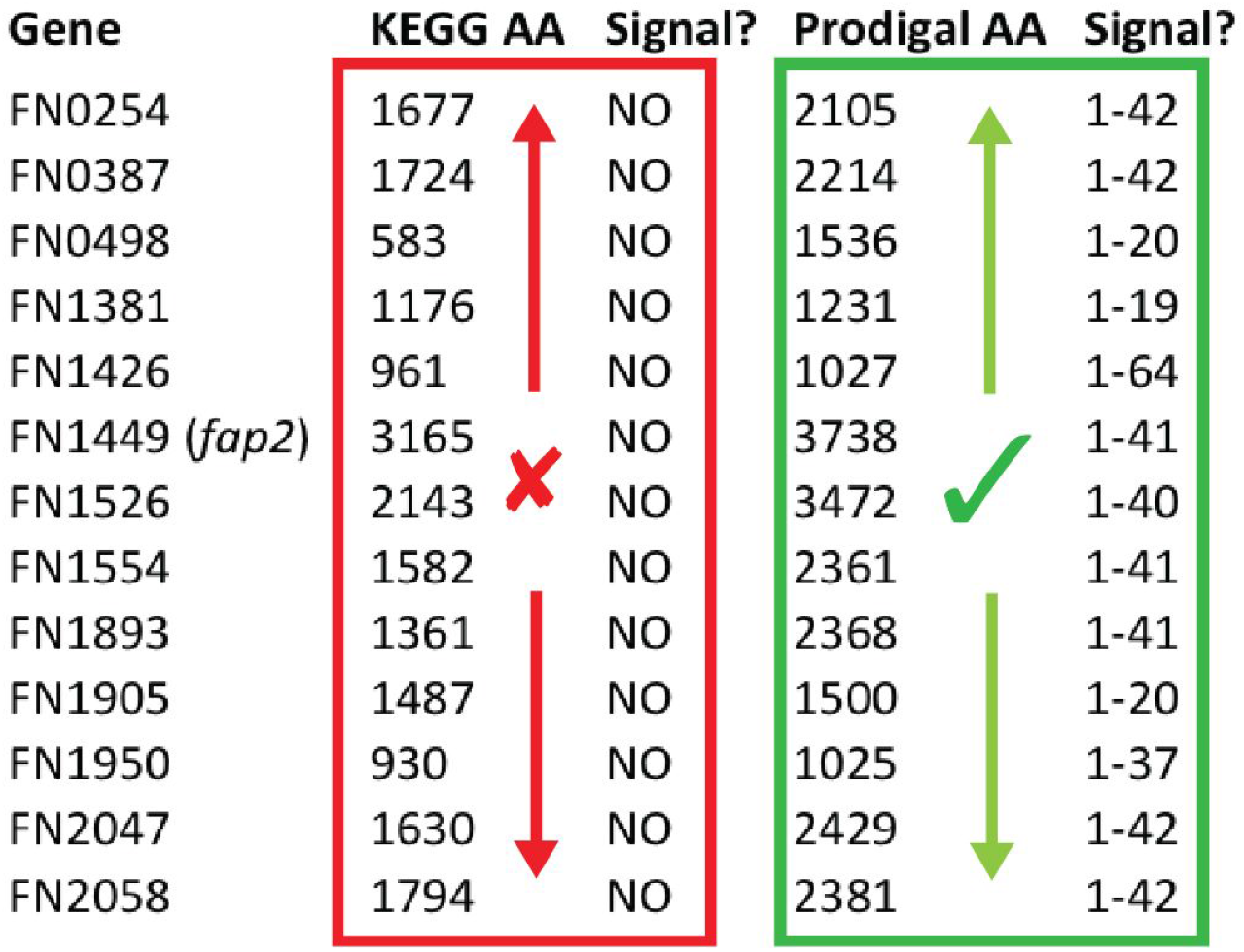
Comparison of T5aSS autotransporter gene annotations in the *F. nucleatum* 25586 genome (GCA_000007325.1) from the KEGG database and our reannotaton using Prodigal. Incorrect gene annotations were not due to an improperly assembled genome, but because of software limitations.

**FIG S2.**
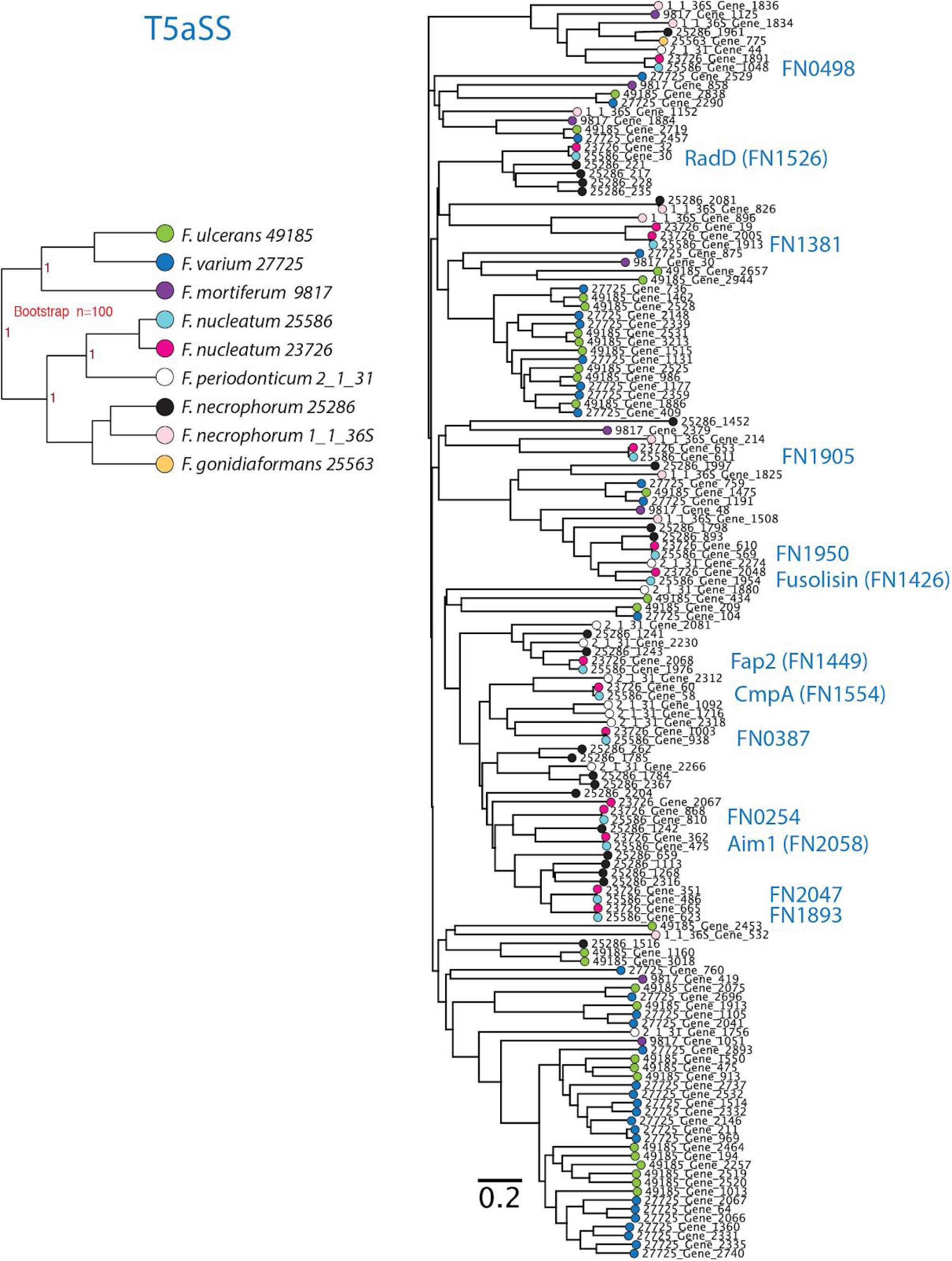
Phylogenetic tree of whole T5aSS autotransporters. Nodes on the tree are colored based on the strain of *Fusobacterium*, and gene names correspond to those found in Table S2 and the FusoPortal database.

**FIG S3.**
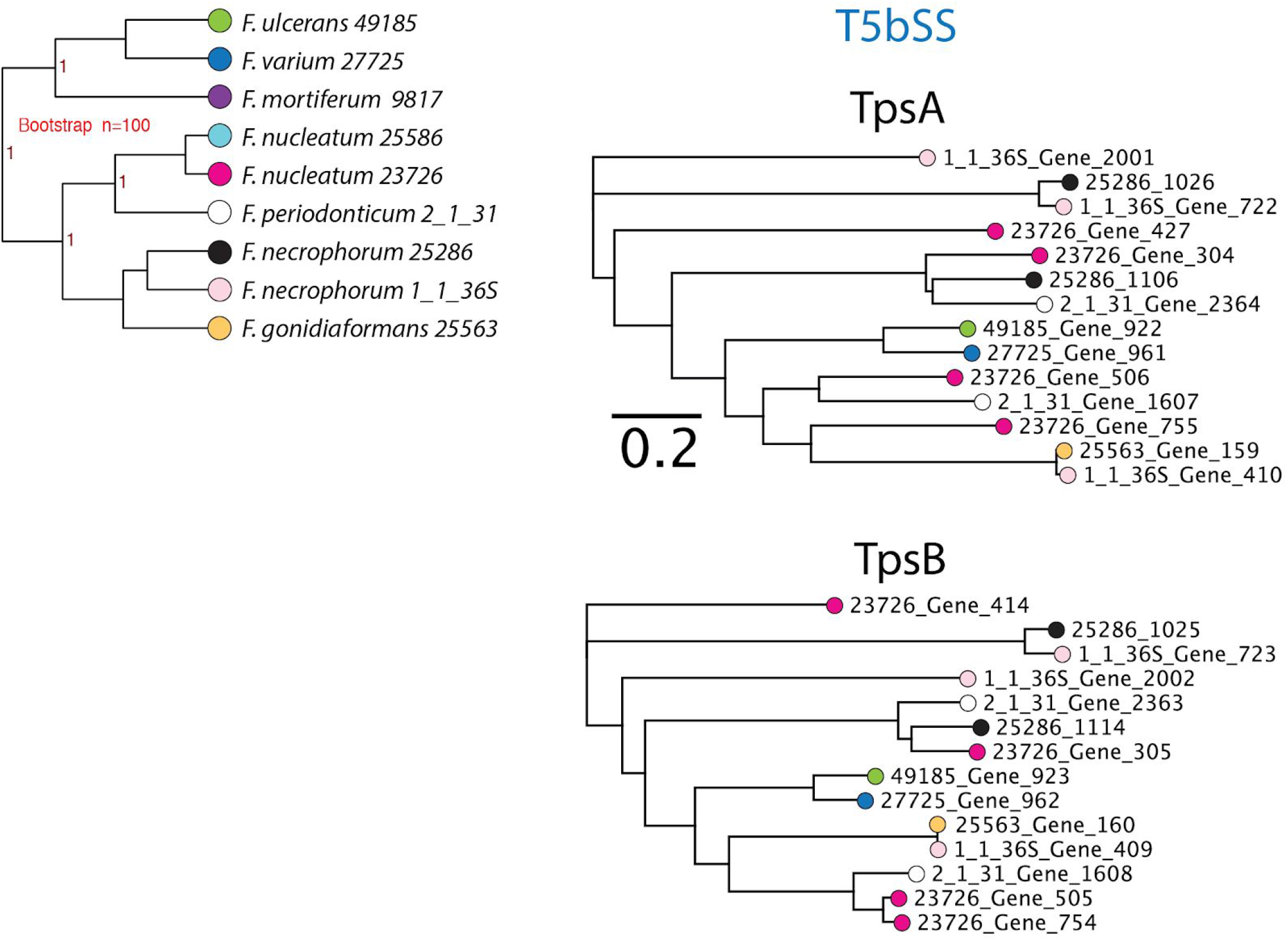
Phylogenetic tree of whole T5bSS two-partner secretion autotransporters. TpsA corresponds to the large, secreted effectors proteins, and TpsB represents the outer membrane embedded β-barrel translocation proteins. Nodes on the tree are colored based on the strain of *Fusobacterium*, and gene names correspond to those found in Table S2 and the FusoPortal database.

**FIG S4.**
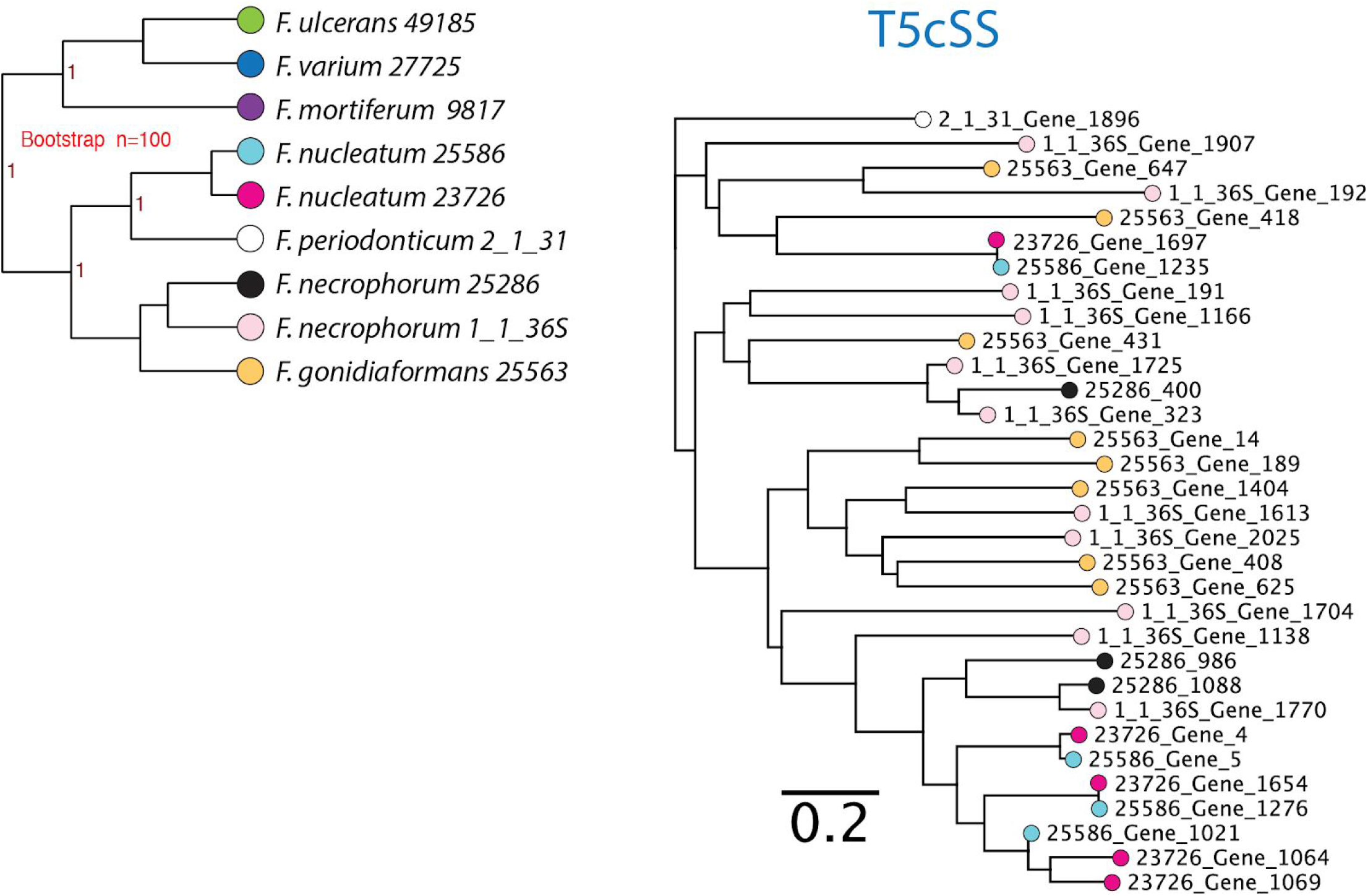
Phylogenetic tree of whole T5cSS trimeric autotransporters. Nodes on the tree are colored based on the strain of *Fusobacterium*, and gene names correspond to those found in Table S2 and the FusoPortal database.

**FIG S5.**
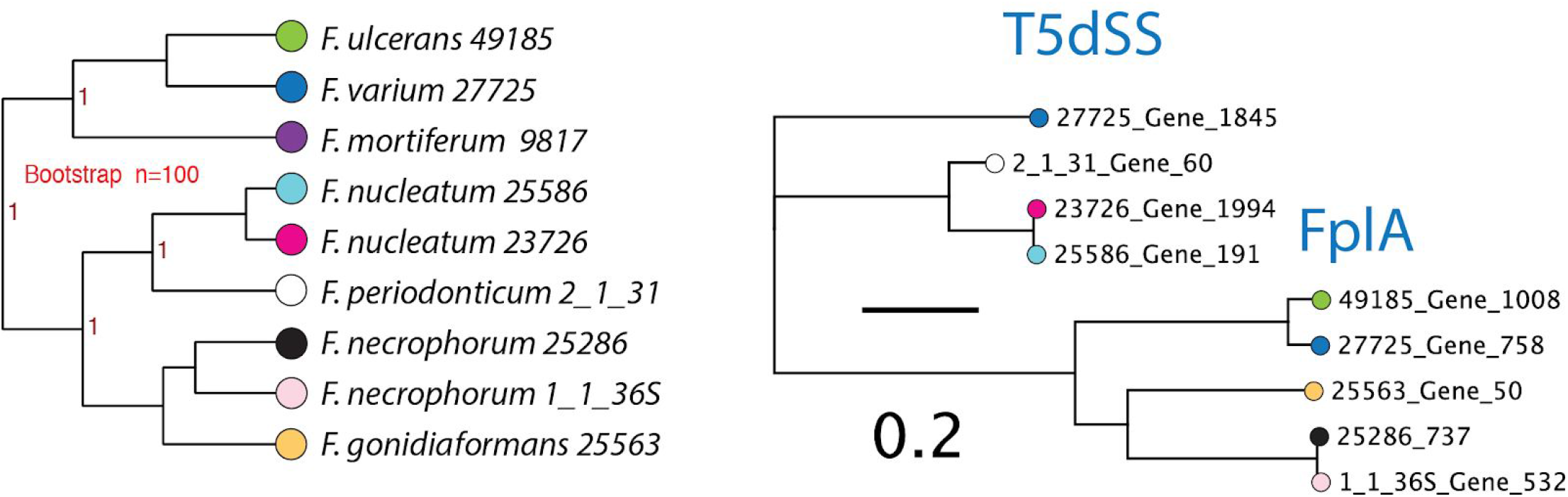
Phylogenetic tree of whole T5dSS phospholipase autotransporters. Nodes on the tree are colored based on the strain of *Fusobacterium*, and gene names correspond to those found in Table S2 and the FusoPortal database.

**FIG S6.**
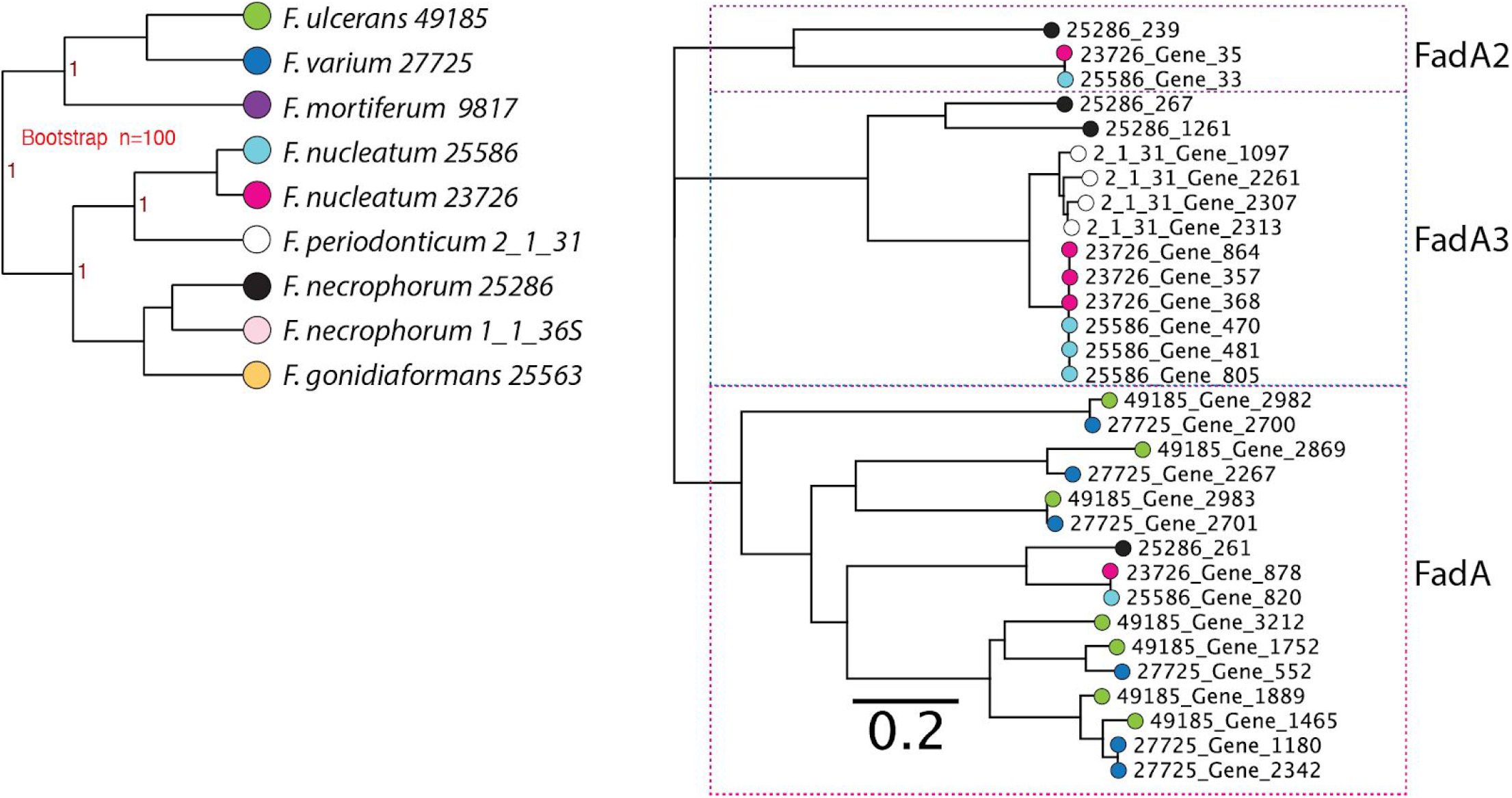
Phylogenetic tree of whole FadA family proteins. Nodes on the tree are colored based on the strain of *Fusobacterium*, and gene names correspond to those found in Table S2 and the FusoPortal database.

**FIG S7.**
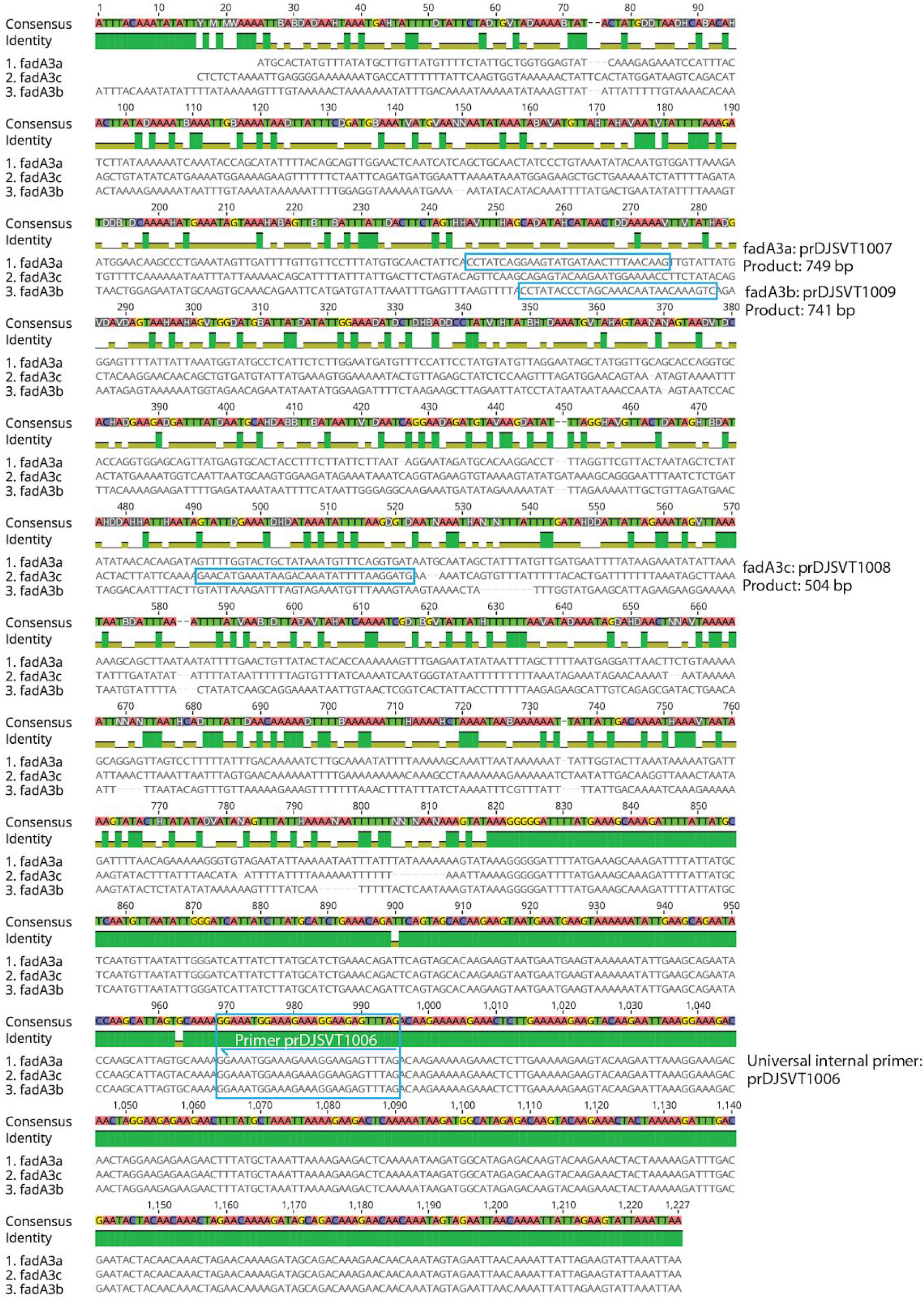
Upstream and coding regions of *fadA3* genes in *F. nucleatum* 23726. A universal reverse primer for gene validation PCR (prDJSVT1006) is paired with a forward primer to produce the indicated bp product to validate these genes are indeed three separate copies in the genome **(Fig. 8D)**.

**FIG S8.**
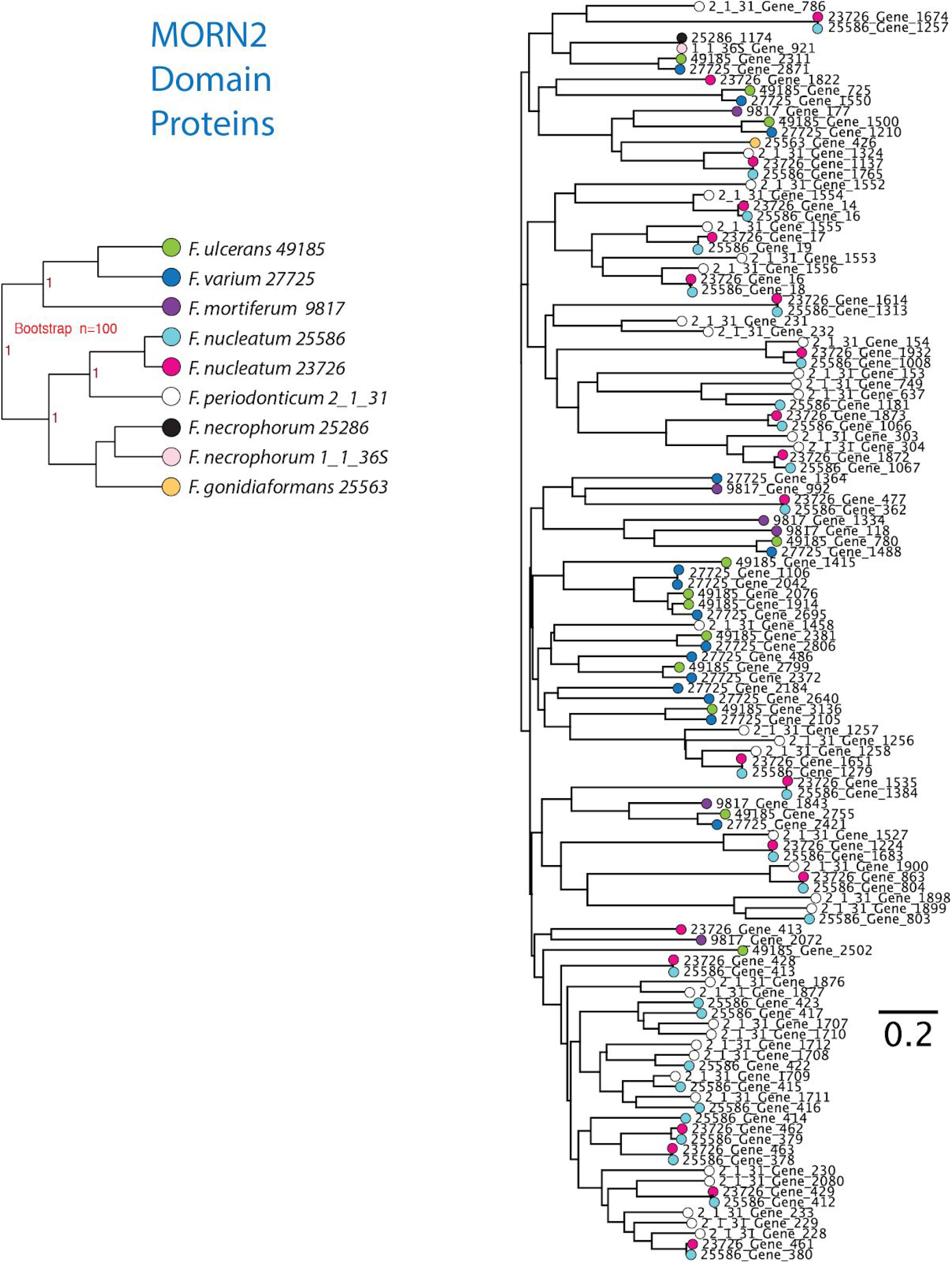
Phylogenetic tree of whole MORN2 domain family proteins. Nodes on the tree are colored based on the strain of *Fusobacterium*, and gene names correspond to those found in Table S2 and the FusoPortal database.

