## Supplementary material for "Reevaluating the *Fusobacterium* Virulence Factor Landscape": Table S1

**Table S2. T5SS autotransporter genes in *F. nucleatum* 25586 and *F. nucleatum* 23726**

### Type 5a Autotransporters (Monomeric autotransporters)

***F. nucleatum* subsp. *nucleatum* ATCC 25586** Blue indicates correction of gene annotation

| FusoPortal Gene | FNXXXX Gene | Assigned Gene Name | Genbank ID° | Uniprot ID | FusoPortal AA | Genbank AA | Uniprot AA | KEGG AA |
|---|---|---|---|---|---|---|---|---|
| Gene_30 | FN1526 | *radD* | NP_602353.1 | Q8RIP5 | **3472** | 2143 | 2143 | 2143 |
| Gene_58 | FN1554 | *cmpA* | NP_602381.1 | Q8RIM1 | **2361** | 1582 | 1582 | 1582 |
| Gene_475 | FN2058 | *aim*1 | NP_602843.1 | Q8RHH1 | **2381** | 1794 | 1794 | 1794 |
| Gene_486 | FN2047 | | NP_602832.1 | Q8RHH7 | **2429** | 1630 | 1630 | 1630 |
| Gene_569 | FN1950 | | NP_602747.1 | Q8R6D6 | **1025** | 930 | 930 | 930 |
| Gene_611 | FN1905 | | NP_602705.1 | Q8RHT9 | **1500** | 1487 | 1487 | 1487 |
| Gene_623 | FN1893 | | NP_602692.1 | Q8RHV1 | **2368** | 1361 | 1361 | 1361 |
| Gene_810 | FN0254 | | NP_603161.1 | Q8RGN7 | **2105** | 1677 | 1677 | 1677 |
| Gene_938 | FN0387 | | NP_603291.1 | Q8RGB7 | **2214** | 1724 | 1724 | 1724 |
| Gene_1048 | FN0498 | | NP_603395.1 | Q8RG21 | **1536** | 583 | 583 | 583 |
| Gene_1913 | FN1381 | | NP_604278.1 | Q8RDW3 | **1231** | 1176 | 1176 | 1176 |
| Gene_1954 | FN1426 | Fusolisin | NP_604320.1 | Q8R608 | **1027** | 961 | 961 | 961 |
| Gene_1976 | FN1449 | *fap2* | NP_604343.1 | Q8RDQ9 | **3440** | 3165 | 3165 | 3165 |

° Genbank ID from previous build under BioProject PRJNA57885. FusoPortal corrected genomes with different accession numbers are under BioProject PRJNA433545.

***F. nucleatum* subsp. *nucleatum* ATCC 23726** Blue indicates correction of gene annotation

| FusoPortal Gene | Corresponding FNXXXX Gene | Assigned Gene Name | Genbank ID$^{\Delta}$ | Uniprot ID | FusoPortal AA | Genbank AA | Uniprot AA | KEGG AA |
|---|---|---|---|---|---|---|---|---|
| Gene_19 | | | EFG95240.1<br>EFG94291.1 | D5RD68<br>D5RFW7 | **1261** | 619<br>649 | 619<br>649 | NA* |
| Gene_32 | FN1526 | *radD* | EFG94737.1 | D5REK4 | 3461 | 3461 | 3461 | |
| Gene_60 | *FN1554* | *cmpA* | EFG94973.1 | D5RDX9 | **2361** | 2072 | 2072 | |
| Gene_351 | FN2047 | | EFG94769.1 | D5REI9 | **2429** | 2010 | 2010 | |
| Gene_362 | FN2058 | *aim*1 | EFG95238.1 | D5RD74 | **2380** | 2336 | 2336 | |
| Gene_610 | FN1950 | | EFG96280.1 | D5RA65 | **1022** | 637 | 637 | |
| Gene_653 | FN1905 | | EFG94596.1 | D5REX7 | 1501 | 1501 | 1501 | |
| Gene_665 | FN1893 | | EFG95107.1 | D5RDJ8 | 2368 | 2368 | 2368 | |
| Gene_868 | FN0254 | | EFG95239.1 | D5RD69 | **2105** | 1752 | 1752 | |

| | | | | | | | |
|---|---|---|---|---|---|---|---|
| Gene_1003 | FN0387 | | EFG95707.1 | D5RBW2 | 2214 | 2214 | 2214 |
| Gene_1891 | FN0498 | | EFG94861.1 | D5RE96 | 1546 | 1546 | 1546 |
| Gene_2005 | FN1381 | | EFG94290.1<br>EFG94291.1 | D5RW8<br>D5RW7 | **1231** | 589<br>649 | 589<br>649 |
| Gene_2048 | FN1426 | Fusolisin | EFG95862.1<br>EFG94287.1 | D5RFX1<br>D5RB84 | **1054** | 700<br>384 | 700<br>384 |
| Gene_2067 | | | EFG95881.1 | D5RBA3 | 2122 | 2122 | 2122 |
| Gene_2068 | Fn1449 | *fap2* | EFG95882.1 | D5RBA4 | 3786 | 3786 | 3786 |

∆ Genbank ID from previous build under BioProject PRJNA31471. FusoPortal corrected genomes with different accession numbers are under BioProject PRJNA433545.
* NA - Not Available - KEGG does not host the previous draft genome of *F. nucleatum* 23726.

### Type 5b Autotransporters (Two partner secretion)

***F. nucleatum*** **subsp.** ***nucleatum*** **ATCC 25586**

**Blue** indicates correction of gene annotation
**Pink** indicates beta-barrel transport gene

| FusoPortal Gene | FNXXXX Gene | Assigned Gene Name | Genbank ID[o] | Uniprot ID | FusoPortal AA | Genbank AA | Uniprot AA | KEGG AA |
|---|---|---|---|---|---|---|---|---|
| Gene_284** | FN0132 | | NP_603039.1 | Q8RGZ3 | **2623** | 2462 | 2462 | 2462 |
| Gene_285 | FN0131 | | NP_603038.1 | Q8RGZ4 | 566 | 566 | 566 | 566 |
| Gene_696 | FN1817 | | NP_602617.1 | Q8RI19 | **2645** | 2806 | 2806 | 2806 |
| Gene_695** | FN1818 | | NP_602618.1 | Q8RI18 | 555 | 555 | 555 | 555 |
| Gene_847[&] | FN0291 | | NP_603198.1 | Q8RGK2 | 1881 | 1881 | 1881 | 1881 |
| Gene_848[&] | FN0292 | | AAL94498.1 | Q8RGK1 | 350 | 350 | 350 | 350 |

[o] Genbank ID from previous build under BioProject PRJNA57885. FusoPortal corrected genomes with different accession numbers are under BioProject PRJNA433545.
[&] Genes are truncated and likely not functional. Gene_847 has a downstream Gene_846 that represents a truncated C-terminus, and Gene_848 (transporter beta barrel) is likely a fragmented version with Gene_849 representing the lost N-terminus.
** Lacks a detectable signal sequences. Previous studies predicted Slipped-Strand Translation could put a signal sequence in frame during phase variation.

***F. nucleatum*** **subsp.** ***nucleatum*** **ATCC 23726**

**Blue** indicates correction of gene annotation
**Pink** indicates beta-barrel transport gene

| FusoPortal Gene | Corresponding FNXXXX Gene | Assigned Gene Name | Genbank ID[∆] | Uniprot ID | FusoPortal AA | Genbank AA | Uniprot AA | KEGG AA |
|---|---|---|---|---|---|---|---|---|
| Gene_304 | FN0132 | *vbaA* | EFG94696.1 | D5REQ7 | 2808 | 2808 | 2808 | NA* |

| | | | | | | | |
|---|---|---|---|---|---|---|---|
| Gene_305 | FN0131 | *vbaB* | EFG94697.1 | D5REQ8 | 596 | 596 | 596 |
| Gene_427 | | *vbbA* | EFG94952.1 | D5RE85 | **2575** | 508 | 508 |
| Gene_414 | | *vbbB* | EFG95313.1 | D5RCX9 | 580 | 580 | 580 |
| Gene_506[&] | | *vbcA* | EFG94580.1 | D5RF52 | 1887 | 1887 | 1887 |
| Gene_505[&] | | *vbcB* | EFG94873.1<br>EFG96294.1 | D5RE06<br>D5RA64 | **534** | 141<br>336 | 141<br>336 |
| Gene_755 | | *vbdA* | | | 2634 | | |
| Gene_754 | | *vbdB* | EFG95529.1<br>EFG96294.1 | D5RCB6<br>D5RA64 | **534** | 152<br>336 | 152<br>336 |

Δ Genbank ID from previous build under BioProject PRJNA31471. FusoPortal corrected genomes with different accession numbers are under BioProject PRJNA433545.

* NA - Not Available - KEGG does not host the previous draft genome of *F. nucleatum* 23726.

[&] Genes are truncated and likely not functional. Gene_847 has a downstream Gene_846 that represents a truncated C-terminus, and Gene_848 (transporter beta barrel) is likely a fragmented version with Gene_849 representing the lost N-terminus.

#### Type 5c Autotransporters (Trimeric autotransporters; YadA like)

***F. nucleatum*** **subsp.** ***nucleatum*** **ATCC 25586** **Blue** indicates correction of gene annotation

| FusoPortal Gene | FNXXXX Gene | Assigned Gene Name | Genbank ID[o] | Uniprot ID | FusoPortal AA | Genbank AA | Uniprot AA | KEGG AA |
|---|---|---|---|---|---|---|---|---|
| Gene_5 | FN1499 | *fvcA* | NP_602326.1 | Q8RIS0 | 479 | 479 | 479 | 479 |
| Gene_1021 | FN0471 | | NP_603368.1 | Q8RG47 | **348** | 340 | 340 | 340 |
| Gene_1235 | FN0691 | *fvcE* | NP_603588.1 | Q8RFK4 | 181 | 181 | 181 | 181 |
| Gene_1276 | FN0735 | *fvcD* | NP_603632.1 | Q8RFG5 | **602** | 617 | 617 | 617 |

o Genbank ID from previous build under BioProject PRJNA57885. FusoPortal corrected genomes with different accession numbers are under BioProject PRJNA433545.

***F. nucleatum*** **subsp.** ***nucleatum*** **ATCC 23726** **Blue** indicates correction of gene annotation

| FusoPortal Gene | Corresponding FNXXXX Gene | Assigned Gene Name | Genbank ID[Δ] | Uniprot ID | FusoPortal AA | Genbank AA | Uniprot AA | KEGG AA |
|---|---|---|---|---|---|---|---|---|
| Gene_4 | FN1499 | *fvcA* | EFG96033.1 | D5RAW6 | 492 | 492 | 492 | NA* |
| Gene_1064 | | *fvcB* | EFG94297.1 | D5RFW5 | **762** | 610 | 610 | |
| Gene_1069 | | *fvcC* | EFG94638.1 | D5RET6 | **615** | 456 | 456 | |
| Gene_1654 | FN0735 | *fvcD* | EFG95777.1 | D5RBK6 | **602** | 617 | 617 | |
| Gene_1697 | FN0691 | *fvcE* | EFG95821.1 | D5RBQ0 | 181 | 181 | 181 | |

∆ Genbank ID from previous build under BioProject PRJNA31471. FusoPortal corrected genomes with different accession numbers are under BioProject PRJNA433545.
* NA - Not Available - KEGG does not host the previous draft genome of *F. nucleatum* 23726.

#### Type 5d Autotransporters (Monomeric phospholipases)

***F. nucleatum* subsp. *nucleatum* ATCC 25586**

| FusoPortal Gene | FNXXXX Gene | Assigned Gene Name | Genbank ID° | Uniprot ID | FusoPortal AA | Genbank AA | Uniprot AA | KEGG AA |
|---|---|---|---|---|---|---|---|---|
| Gene_191 | FN1704 | *fplA* | NP_602520.a | Q8R6F6 | 760 | 760 | 760 | 760 |

° Genbank ID from previous build under BioProject PRJNA57885. FusoPortal corrected genomes with different accession numbers are under BioProject PRJNA433545.

***F. nucleatum* subsp. *nucleatum* ATCC 23726**

| FusoPortal Gene | Corresponding FNXXXX Gene | Assigned Gene Name | Genbank ID∆ | Uniprot ID | FusoPortal AA | Genbank AA | Uniprot AA | KEGG AA |
|---|---|---|---|---|---|---|---|---|
| Gene_199 | FN1704 | *fplA* | EFG94447.1 | D5RFI3 | 760 | 760 | 760 | NA* |

∆ Genbank ID from previous build under BioProject PRJNA31471. FusoPortal corrected genomes with different accession numbers are under BioProject PRJNA433545.
* NA - Not Available - KEGG does not host the previous draft genome of *F. nucleatum* 23726.
